# Gene signature of circulating platelet-bound neutrophils is associated with poor prognosis in cancer patients

**DOI:** 10.1101/2021.08.26.457803

**Authors:** Pacôme Lecot, Maude Ardin, Sébastien Dussurgey, Vincent Alcazer, Lyvia Moudombi, Margaux Hubert, Aurélie Swalduz, Hector Hernandez-Vargas, Alain Viari, Christophe Caux, Marie-Cécile Michallet

## Abstract

Beyond their critical role in hemostasis, platelets physically interact with neutrophils to form neutrophil-platelet aggregates (NPAs), enhancing neutrophil effector functions during inflammation. NPAs may also promote disease worsening in various inflammatory diseases. However, characterization of NPAs in cancer remains totally unexplored. Using ImageStream®X (ISX) imaging flow cytometer, we were not only allowed able to detect CD15^+^ CD14^−^ CD36^+^ ITGA2B^+^ NPAs in both healthy donors’ (HDs) and cancer patients’ bloods, but we also showed that NPAs result from the binding of platelets preferentially to low-density neutrophils (LDNs) as opposed to normal-density neutrophils (NDNs). By re-analyzing two independent public scRNAseq data of whole blood leukocytes from cancer patients and HDs, we could identify a subset of neutrophils with high platelet gene expression that may correspond to NPAs. Moreover, we showed that cancer patients’ derived NPAs possessed a distinct molecular signature compared with the other neutrophil subsets, independently of platelet genes. Gene ontology (GO) term enrichment analysis of this NPAs-associated neutrophil transcriptomic signature revealed a significant enrichment of neutrophil degranulation, chemotaxis and trans-endothelial migration GO terms. Lastly, using The Cancer Genome Atlas (TCGA), we could show by multivariate Cox analysis that the NPAs-associated neutrophil transcriptomic signature was associated with a worse patient prognosis in several cancer types. These results suggest that neutrophils from NPAs are systemically primed by platelets empowering them with cancer progression capacities once at tumor site. NPAs may therefore hold clinical utility as novel non-invasive blood prognostic biomarker in cancer patients with solid tumors.

**Novelty and Impact:** Platelets physically interact with peripheral blood neutrophils to form neutrophil-platelet aggregates (NPAs), known to promote disease worsening in various inflammatory diseases. However, characterization of NPAs in cancer remains totally unexplored. We showed that NPAs-associated neutrophils were a yet-unreported unique subset of circulating neutrophils associated with a worse patient prognosis in several cancer types. NPAs may hold clinical utility as novel non-invasive blood prognostic biomarker in cancer patients with solid tumors.

## Background

Although, some recent evidences suggested that tumor-associated neutrophils (TANs) could display anti-tumor properties [1–3], a large body of evidence demonstrated that TANs could be pro-tumoral and promote metastasis [4–6]. Pro-tumor TANs can assist metastasis by blunting anti-tumor T-cell responses [4], and stimulating proliferation and invasiveness of metastatic tumor cells [5,6]. Accumulating evidence in mice and humans showed that blood neutrophils may also contribute to cancer progression. Numerous retrospective analysis across multiple different cancer types showed that an elevation in the absolute count of circulating neutrophils together with a decrease in the absolute count of circulating lymphocytes, both accounting for a high neutrophils-to-lymphocytes ratio (NLR) on blood, was associated with a worse prognosis in patients [7–10]. More recently, studies have shown that a subset of circulating neutrophils, named low-density neutrophils (LDN) expanded in cancer patient’s blood and was associated with worse prognosis, especially in non-small-cell lung cancer (NSCLC) and head and neck cancer (HNC) [11,12]. Moreover, LDN was found to display features of pro-tumor neutrophils, such as increased T-cell suppressive functions, as opposed to normal-density neutrophils (NDN) [11,13]. Nevertheless, LDN remains a heterogeneous population of cells including mature and immature neutrophils [13,14]. A subset of circulating LDN expressing the LOX-1 scavenger receptor was reported to display higher T-cell suppressive functions as compared to LOX-1^−^ LDN [13]. LOX-1^+^ T-cell suppressive neutrophils were recently reported to infiltrate lung tumors and were associated with a worse patient survival [15]. Besides studies in humans, recent evidence in mice demonstrated that the pre-programming of neutrophils in the periphery was required for acquisition of their tumor-promoting functions once at tumor site [16,17] supporting the idea that tumor-promoting TANs derived from a distinct subset of neutrophils pre-existing in the periphery.

Beyond their critical role in hemostasis, platelets can bind to circulating leukocytes forming leukocyte-platelet aggregates, enhancing their effector functions. For instance, platelets have been recently shown to interact with peripheral blood monocytes to activate antigen cross-presentation [18]. In sepsis, in contrast to free neutrophils, NPAs (neutrophil-platelet aggregates) produce higher amount of neutrophil extracellular traps (NETs) [19–21] and display increased phagocytic functions [22–24]. Furthermore, NPAs display enhanced trans-endothelial migration capacity [25,26] therefore promoting neutrophil infiltration to site of injury and fueling pathological inflammation in various diseases [21,27–29]. However, to our knowledge, the characterization of NPAs in cancer patients remains unexplored.

In this study, we showed that neutrophils from NPAs represent a unique subset of activated neutrophils in cancer patients by combining ImageStream®X (ISX) imaging flow cytometry and analysis of public single cell transcriptomic data of human circulating neutrophils. In particular, we showed that LDN as opposed to NDN were preferentially involved in NPA formation. Finally, we could generate a specific gene signature of neutrophils in NPAs that we found associated with a worse prognosis in pancreatic adenocarcinoma patients and liver hepatocellular carcinoma patients.

## Materials and Methods

### Human cohorts

Whole blood from a cohort of 6 metastatic NSCLC patients (Table S1) who gave informed consent at the Léon Bérard Hospital (France), was collected in EDTA-coated tubes. As an age and sex-matched control cohort, whole blood from healthy donors (HD) was collected via the Etablissement Français du Sang (Table S1).

### Preparation of whole blood, NDNs and LDNs-enriched fractions for multi-spectral Imaging Flow Cytometry (ImageStream®X, ISX)

Blood from NSCLC patients or HD was withdrawn in EDTA-coated vacutainers 1 hour prior to the processing. Two tubes of 3 ml of blood were centrifuged at 800 rpm (no brake) for 15 min at room temperature (RT) to separate plasma from leukocyte-enriched red pellet. Platelet-rich plasma was discarded for both tubes. One out of the two tubes was used for whole blood staining while the other served for the separation of LDNs from NDNs. For such separation, leukocyte-enriched red pellet was re-suspended in 6 ml of PBS_1X_ (5 mM EDTA) and then layered on the top of 3 ml of Ficoll (CMSMSL01-01, EUROBIO), before being centrifuged at 1800 rpm for 20 min at RT (acceleration 5 and brake 1). LDNs-enriched PBMC ring and NDNs-enriched red pellet were washed with PBS_1X_, and pellets were re-suspended in BD Pharm Lyse™ (555899) according to manufacturer’s protocol to eliminate red blood cells. In parallel, whole blood leukocyte-enriched red pellet was also re-suspended in BD Pharm Lyse™ for the same purpose. After incubation, red blood cells (RBC)-lysed leukocytes were washed twice with PBS_1X_ (5 mM EDTA) were re-suspended at a final concentration of 20.10^6^ cells/ml prior to be stained. Samples were acquired on ImageStream®X (ISX) no more than 5 days following cell staining and fixation.

Two millions cells per sample were stained for 30 min in the dark at 4°C in 100 μl staining buffer (PBS_1X_, 2 % SVF, 5 mM EDTA) with antibodies directed against CD15 (561584, Clone HI98), CD14 (562335, MφP9), CD41a (559777, HIP8) (all from BD), and CD36 (130-110-739, REA760) and CD62P (130-105-714, REA389) from Miltenyi Biotec. Cells were washed twice with PBS_1X_ (5 mM EDTA, 2% fetal bovine serum (FBS)) before being fixed in 2% formaldehyde solution (252549, Sigma-Aldrich). After incubation at 4°C, cells were washed twice with PBS_1X_ (5 mM EDTA, 2% FBS) and re-suspended in a final volume of 180 μl PBS_1X_ (5 mM EDTA, 2% FBS).

NPAs were imaged with ImageStream®X (ISX) imaging flow cytometer (Amnis Corporation-Luminex, Seattle, WA) using 405, 488, 561, and 642 lasers and the 40X objective. Brightfield provided morphological and structural details of the cell. At least 300 000 RBC-lysed blood cells from NSCLC patients or HDs separately, excluding debris and free-platelet with low area, were collected for each sample. Since flow-speed could influence the proportion of NPAs [30], RBC-lysed blood cells were acquired with a flow-speed between 500 to 1000 events/second for each sample. Data were analyzed using IDEAS image analysis software (Amnis Corporation, Seattle, WA). A specific gating strategy, with multiple filtering steps was set up to identify non-coincidental viable NPAs. Such strategy aimed first at retaining only focused cells based on Gradient RMS values for each event. Remaining debris and free platelets with low area were discarded. Dead cells positive for Zombie NIR (Biolegend) viability marker were then eliminated. Neutrophils were selected as cells with a high expression of CD15 and a low expression of CD14 markers. Low area event on CD15 object mask were eliminated to exclude CD15^+^ debris or CD15^+^ insufficient quality staining for an accurate estimation of true NPAs. Platelet-related events were then selected by gating on events positive for both ITGA2B and CD36 cell-surface markers. A second filter was applied to narrow down the selection of platelet-related events by only considering the ones with high ITGA2B intensity on ITGA2B-related component with the largest area to remove residual platelet debris. To only retain non-coincidental NPAs, two distinct masks delimiting neutrophil-related CD15 stained-area and platelet-related ITGA2B stained-area were created. Non-coincidental NPAs were events for which neutrophil and platelet-associated masks were overlapping with at least one pixel, and confirmed by image examination of random samples. Only the most-focused platelet-related events were considered by selecting events with high gradient RMS of ITGA2B marker on platelet area. Frequencies of NPAs were calculated on at least 100 non-coincidental NPAs for each NSCLC patients and HDs.

### May-Grünwald-Giemsa staining

A blood smear from a drop of NSCLC patients’ whole blood was performed on a glass slide. Blood smear was dried up for 10/15 min at RT. May-Grünwald staining was layered on dried blood smear using Hema-Tek Stain Pak Kits designed for the Bayer Hema-Tek 2000 Slide Stainer. Pumps deliver fresh reagents (Stain, Buffer and Rinse solution) in precise volume. Slide was then taken out and completely dried at RT before being covered glasses and liquid mounting media using Tissue-Tek Coverslipping Film. Slides are immediately available for being digitized on Pannoramic scan II slide scanner by 3DHistech (x40, Z Stack, Multilayer Mode 5*0,2um) and being visualized using case viewer software.

### Bioinformatic analysis of public scRNAseq data of human whole blood leukocytes

#### ScRNAseq data of NSCLC patients’ whole blood leukocytes (GSE127465)

ScRNAseq data of 6 NSCLC patients’ whole blood leukocytes were downloaded from GEO as normalized count per million (CPM) gene expression matrix. Briefly, pre-processing of scRNAseq data performed by the authors [31] including the removal of: i) dead cells identified by a mitochondrial gene content > 20% among total gene transcribed ; ii) cells with poor quality transcriptome, as being cells with less than 300 counts ; iii) doublet cells performed with Scrublet [32]. We next performed analysis of CPM gene expression matrix from 6 NSCLC patients’ whole blood leukocytes by using Seurat R package (version 3.1.1). We first removed genes that were not expressed in at least 3 cells and filtered-out cells with less than 200 different transcribed-genes (Seurat default parameters). Matrix was log-transformed and scaled prior to principal component analysis (PCA) with ScaleData Seurat function. PCA was performed on the 2000 most-variable genes, using RunPCA function. We used JackStraw function to determine the statistical significance of PCA scores that led us to retain the first 9 principal components (PCs). We next define the number of clusters by using FindNeighbors and FindClusters functions that take into account the first 9 PCAs and a resolution of 0.66, leading to 11 clusters. We ran the uniform manifold approximation and projection (UMAP) dimensional reduction technique, using RunUMAP function, and then display 2D UMAP projections. We used the authors major cell type annotations [31] based on the validated leukocyte gene signature matrix (LM22) [33].

#### ScRNAseq data of healthy donors’ whole blood leukocytes (GSE145230)

ScRNAseq data of 3 human males’ and 4 human females’ whole blood leukocytes were downloaded from GEO as normalized count per million (CPM) gene expression matrix. Pre-processing of scRNAseq data was performed identically as described above for GSE127465, except that doublet cells were removed based on UMI number and % of mitochondrial gene expression [34]. We next define the number of clusters by using FindNeighbors and FindClusters functions that take into account the first 19 PCAs and a resolution of 0.6, leading to 22 clusters. Cells were annotated by combining Clustifyr (version 1.0.0) [35] (using reference matrices from Seurat CBMC, Zilionis Blood datasets [31], and MCP-counter genes list [36]) with SingleR (version 1.2.4) [37] annotation tools (Human Primary Cell Atlas used as reference).

#### Differential gene expression analysis

From Seurat object generated from GSE127465 CPM gene expression matrix, differential gene expression analysis between 2 distinct clusters was performed using FindAllMarkers function from Seurat R package. Only genes detected in at least 25% of cells and with a positive average log_2_ fold change of 0.25 were retained, but no cutoff value for the adjusted p-value was used. Union function from dplyr R package (version 1.0.2) was used to get the union of upregulated genes (Figure 4.B ; Table S4) from multiple differential gene expression analysis. Setdiff function was used to get the exclusive list of upregulated genes of a given cluster by opposition to another one.

#### Single sample Gene Set Enrichment Analysis (ssGSEA) on bulk transcriptomic data

SsGSEA from GSVA R package (version 1.32.0) was performed on public microarray data of *in vitro* stimulated and unstimulated neutrophils (GSE15139 and GSE49757). For both data sets, control probes, and probes with ambiguous gene symbol were removed. For probes matching to identical genes, mean expression value per gene was calculated on probes matching to that gene to retain only one gene expression value per gene. SsGSEA of the Raghavachari platelet signature and the Ponzetta neutrophil signature (Table S2) was performed on each replicate of each group using GSVA R package. All ssGSEA scores were bar-plotted with ggplot2 R package (version 3.2.1).

#### Single sample Gene Set Enrichment Analysis (ssGSEA) on scRNAseq data

ssGSEA from R escape package (version 0.99.9) was used for calculating an enrichment score for Raghavachari platelet signature and Ponzetta neutrophils signature (Table S2) of all individual cells. Only cells with at least 1 count per gene present in Raghavachari or Ponzetta signatures were considered for calculating the enrichment score. Resulting ssGSEA scores obtained for each cluster was plotted using ggviolin function from ggpubr R package (version 0.4.0).

#### Venn diagrams

Venn diagrams were generated by using the Venn and compute.Venn functions from the Vennerable package (version 3.1.0.9000).

#### MCP counter on scRNAseq data

Normalized CPM expression matrix for all cells was generated from Seurat object. MCP counter scores (mean expression of all genes for a given signature (in log_2_ TPM +1)) of the “16_gene_NPA_signature” (Table S5) were calculated for each individual cells using MCPcounter.estimate function from MCPcounter package (version 1.2.0). Comparison of mean MCP score between NPA cluster (Neu 5 PPBP^high^ PF4^high^ NRGN^high^ cells) and each individual neutrophil clusters was performed with the Stat_compare_means function from the ggpubr package (version 0.4.0) choosing non-parametric Wilcoxon rank-sum (Mann–Whitney) statistical test. Results were presented through violin plots that were generated with ggplot2 R package (version 3.3.2).

### Survival analysis

TCGA pan-cancer RSEM normalized log2 transformed gene expression data were downloaded from UCSC Xena Browser and log2 +1 transformed. Updated clinical data with survival endpoints were retrieved from TCGA-CDR paper (Cell 2018, https://doi.org/10.1016/j.cell.2018.02.052).

Immune signatures (including the “NPA_12”) were calculated as the mean expression of each individual gene. Patients were then stratified into three groups based on each signature’s tercile values.

Survival analysis was performed using the survminer and survival R packages. For each cancer type, survival curves were estimated with Kaplan Meier and compared with the log-rank test. Multivariate analyses considering age and tumor stage as cofounding factors were then performed using a cox model with p-value correction using the Benjamini-Hochberg method.

## Results

### Identification of NPAs in peripheral blood of cancer patients and healthy donors (HDs)

NPAs from NSCLC patients and HDs’ whole blood were identified as CD15^+^ CD14^−^ neutrophils that were double positive for platelet glycoprotein 4 (CD36) and integrin alpha-IIb (ITGA2B, also known as CD41a) by using ImageStream®X (ISX) imaging flow cytometer (Fig. 1A-B). We could also easily identify NPAs in unprocessed NSCLC patients’ whole blood smear through May–Grünwald–Giemsa staining (Fig. 1C). Because the frequency of NPAs was shown to be increased in the blood of patients with various pathologies as compared to HDs [38–40], we next quantified by ISX the frequency of NPAs among neutrophils between NSCLC patients and HDs with the same gating strategy. Although the frequency of NPA between NSCLC patients (mean frequency = 1.48%) and HDs (mean frequency = 1.16%), was roughly the same, with no statistically significant difference (p value = 0.64), one NSCLC patient displayed more than a three-fold increase of NPA proportion (mean frequency = 4.59%) compared to the mean frequency of NPA in NSCLC patients and HDs (Fig. 1D). Interestingly, among all 6 NSCLC patients analyzed, this patient with high percentage of NPAs also displayed by far the highest NLR value (34.56 as opposed to 7.21 for the others), mostly explained by a higher absolute neutrophil count (Table S1), and the highest tumor burden (7 vs 2 metastatic sites in the rest of the cohort) (Table S1).

**Figure 1.**
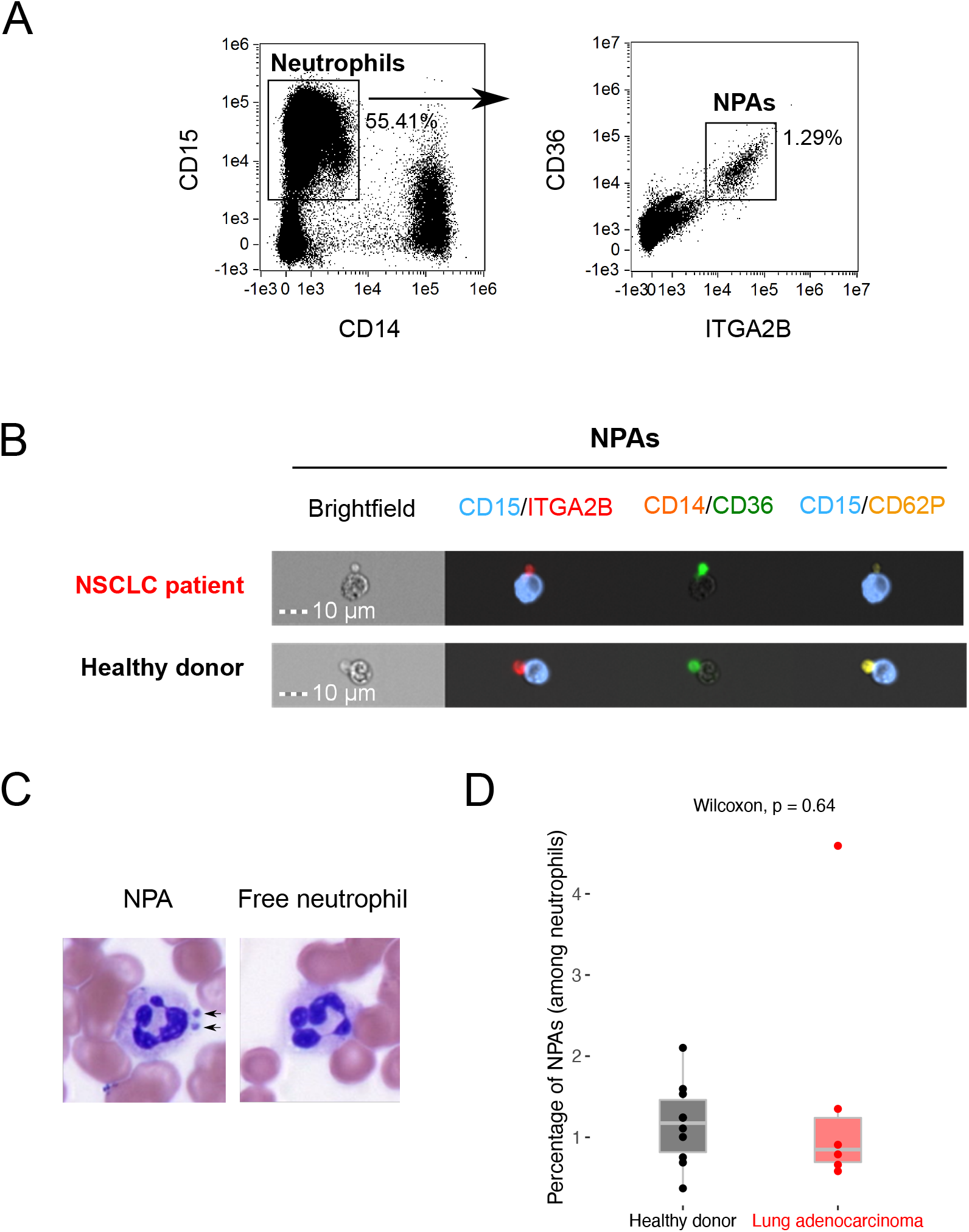
Identification of NPAs in both NSCLC patients and healthy donors’ peripheral blood. *(**A**)* Gating strategy on ISX to identify CD36^+^ ITGA2B^+^ NPAs from CD15^+^ CD14^−^ neutrophils in NSCLC patients’ peripheral whole blood (one representative data of n= 6 NSCLC patients’ blood). *(**B**)* Visualization of CD15^+^ CD14^−^ CD36^+^ ITGA2B^+^ NPAs by ISX in NSCLC patients and HDs’ peripheral blood. Images were taken with X40 objective. CD15: Neutrophil marker; CD14: Monocyte marker; ITGA2B, CD36, CD62P: Platelet markers. *(**C**)* May-Grünwald-Giemsa coloration on NSCLC patient’s whole blood smear. X100 magnification. Dark arrows show the presence of platelets aggregated to neutrophil (NPAs). Neutrophil is evidenced by segmented nucleus in dark purple. Free neutrophil refers to neutrophil free of platelets. *(**D**)* Box plot representing the frequency of non-coincidental CD15^+^ CD14^−^ CD36^+^ ITGA2B^+^ NPAs (among all neutrophils) in whole blood of NSCLC (n=6) patients and HDs (n=10), determined by ISX. Difference in terms of frequency of NPAs between NSCLC patients and HDs was assessed by the Wilcoxon rank-sum (Mann–Whitney) statistical test.

### Platelets preferentially bind to low-density neutrophils (LDNs), rather than normal-density neutrophils (NDNs) in cancer patients’ blood

Given the well-known detrimental role of LDNs in cancer [11–14,41,42], we next analyzed a potential link between LDNs and NPAs.

We first performed a single sample Gene Set Enrichment Analysis (ssGSEA) on publicly available transcriptomic data of bulk LDNs and bulk NDNs from NSCLC and Head and Neck Cancer (HNC) patients’ blood (GSE79404). We showed that the published platelet-specific gene signature “Raghavachari” [43] (Table S2) was significantly (p-value = 0.0022) more enriched in LDNs than in NDNs (Fig. 2A).

**Figure 2.**
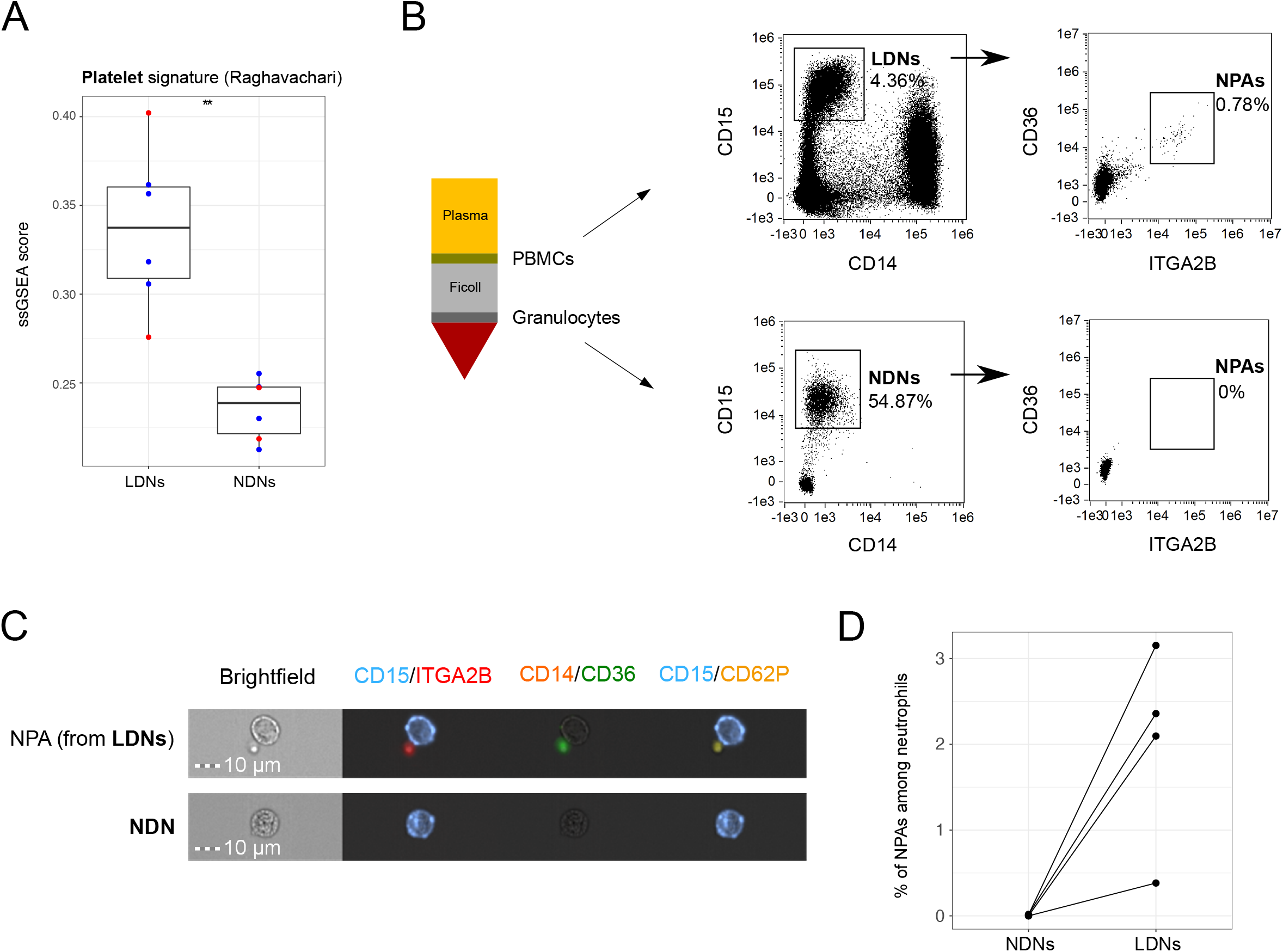
NPAs define a subset of LDNs not present in NDNs. *(**A**)* Box plot representing ssGSEA enrichment scores of the Raghavachari platelet signature (Table S2) across LDNs and patients’ matched NDNs from four HNC (blue dots) and two NSCLC (red dots) cancer patients (GSE79404). Differential enrichment of platelet signature between LDN and NDN groups was assessed by the Wilcoxon rank-sum (Mann–Whitney) statistical test. P-value was displayed on graphs, ** p ≤ 0.01. *(**B**)* Schematic representation of Ficoll density gradient centrifugation of human whole blood showing the separation of the low-density fraction (containing PBMCs) from the normal-density fraction (containing mostly granulocytes). Both fractions from NSCLC patients’ blood were analyzed by ISX, to identify CD15^+^ CD14^−^ LDNs (in the low-density fraction) and CD15^+^ CD14^−^ NDNs (in the normal-density fraction). From either LDNs or NDNs gates, all CD36^+^ ITGA2B^+^ events were gated as “NPAs”. *(**C**)* Visualization by ISX in NSCLC patients of non-coincidental CD36^+^ ITGA2B^+^ NPAs from LDN gate in the PBMC fraction and NDN gate in the granulocyte fraction. Images were taken with X40 objective. CD15: Neutrophil marker; CD14: Monocyte marker; ITGA2B, CD36, CD62P: Platelet markers. *(**D**)* Dot plot reflecting the frequencies of non-coincidental CD15^+^ CD14^−^ CD36^+^ ITGA2B^+^ NPAs among NDNs and LDNs determined by ISX, across n=4 NSCLC patients.

We next isolated LDNs and NDNs (Fig. 2B). Using ISX, we could visualize CD15^+^ CD14^−^ CD36^+^ ITGA2B^+^ NPAs from LDNs but not from NDNs (Fig. 2B-C). NPAs represented around 2% of LDNs but were negligible in the NDN population (Fig. 2B-D). Collectively, these results demonstrate that platelet are preferentially bound to LDNs rather than NDNs.

### Identification of NPAs in public scRNAseq datasets from whole blood leukocytes in both cancer patients and HDs

We next aimed at characterizing NPAs from cancer patients at the transcriptomic level. Indeed, given the small size of platelets, we hypothesized that NPAs could be isolated as a single cell (Fig. 1B), just like free neutrophil, and therefore be present in scRNAseq data. For this purpose, we took advantage of a recently published scRNAseq dataset of NSCLC patients’ whole blood leukocytes (GSE127465) [31]. Therefore, for our analysis we used normalized counts tables and kept author’s major cell type annotations to identify blood cell populations. We could re-cluster all leukocytes (Fig. 3A) and distinguished 6 clusters of neutrophils (named Neu 1, Neu 2, Neu 3, Neu 4, Neu 5 and Neu 6 (Fig. 3B). Interestingly, we observed that Neu 5 cluster seemed to be distinct from the other neutrophil clusters, through its close proximity with the platelet cluster (Fig. 3B). Using a published neutrophil-specific gene signature “Ponzetta” (Table S2), we showed that cells from Neu 5 had a ssGSEA enrichment score (median ssGSEA score = 0.21) similar to the other neutrophil clusters (median ssGSEA score = 0.24), and greater than cells from the other cell types clusters (median ssGSEA score = -0.15) (Fig. 3C), validating that cells from Neu 5 are neutrophils. This was supported by the highest expression level of neutrophil-specific genes (*CSF3R, CXCR2* and *FCGR3B*) in all six clusters of neutrophils, including Neu 5 cluster (Fig. 3E). We then evaluated the ssGSEA enrichment score of the published platelet-specific gene signature “Raghavachari” (Table S2) (33) across all clusters identified. Platelet cluster was by far the most enriched cell type for the platelet-specific gene signature (median ssGSEA score = 0.25 versus -0.28 in the other clusters), validating the specificity of this signature (Fig. 3D). We found that a fraction of the Neu 5 neutrophils as well as of monocytes were more enriched for the platelet-specific gene signature than any other of the non-platelet cell types (Fig. 3D). Looking at the gene level, *PF4*, *PPBP* and *NRGN* (Table S3) platelet genes were in the 5 most-differentially upregulated genes (Table S4) able to discriminate the Neu 5 cluster from the other neutrophils and immune cell clusters (Fig. 3F). In conclusion, the Neu 5 cluster of platelet gene-expressing neutrophils seems to correspond to NPAs.

**Figure 3.**
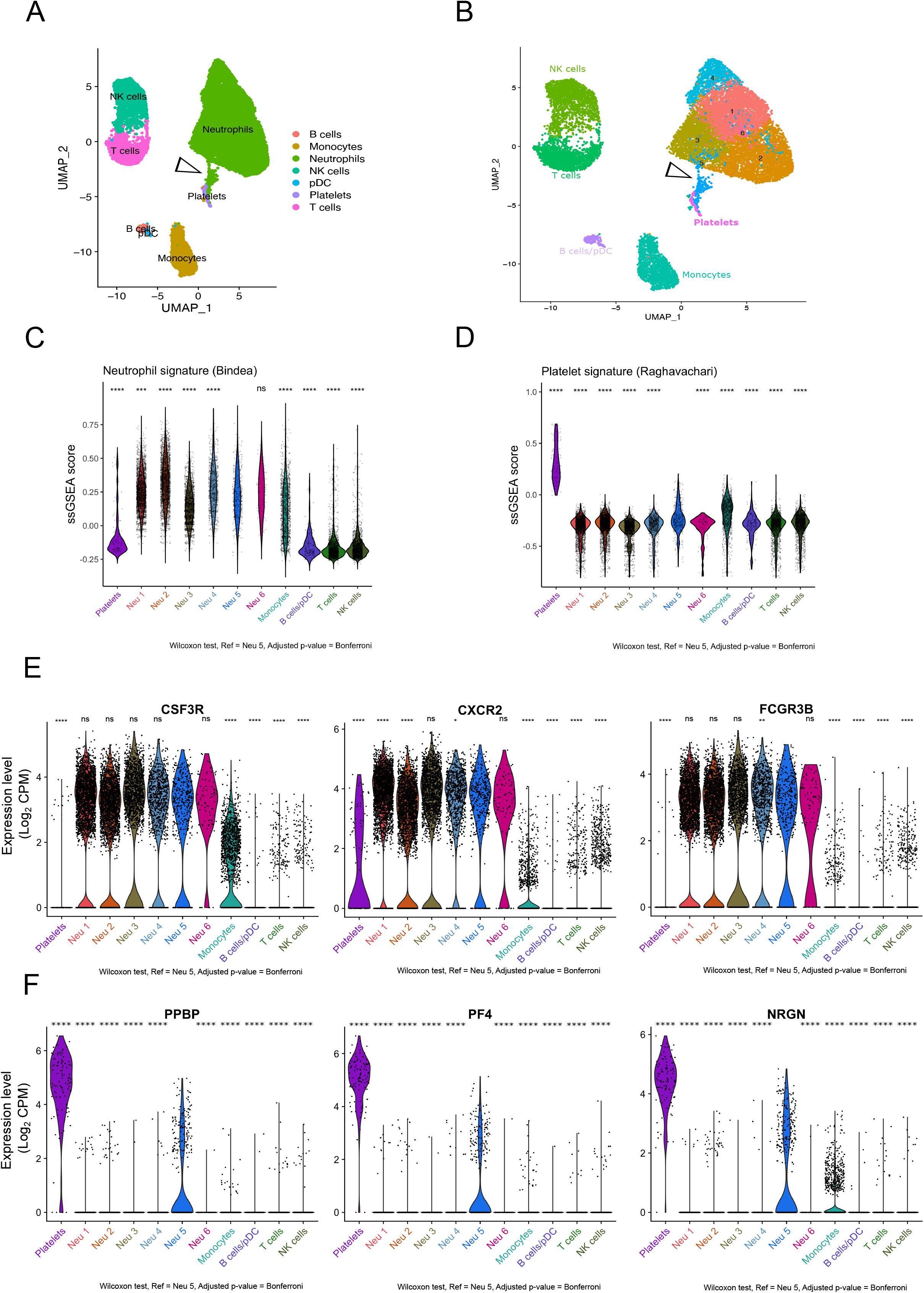
Identification of a cluster of neutrophils expressing platelet genes in public scRNAseq data from NSCLC patients’ peripheral blood leukocytes. *(**A-B**)* Two dimensional-UMAP representation of re-clustered NSCLC patients’ whole blood leukocytes from scRNAseq data (GSE127465). *(**A**)* Major immune cell types were labeled using Zilionis et al. annotation. ***(B)*** Each of the 11 clusters defined by the analysis were labeled based on the type of immune cell. The 6 clusters of neutrophils were annotated as follow: Neu 1, Neu 2, Neu 3, Neu 4, Neu 5 and Neu 6. Dark arrows show Neu 5 neutrophil cluster of interest displaying high expression of platelet genes (see differential gene expression analysis Table S3). *(**C**)* Violin plot representing ssGSEA enrichment score per cell of the Ponzetta neutrophil signature (Table S2) across all clusters of blood immune cells. *(**D**)* Violin plot representing ssGSEA enrichment score per cell of the Raghavachari platelet signature (Table S2) across all clusters of blood immune cells. *(**E**)* Violin plots representing the log_2_ gene expression (count per million, CPM) of neutrophil-specific genes (*CXCR2*, *CSF3R* and *FCGR3B*) (Table S2) across all clusters of blood immune cells. *(**F**)* Violin plots representing the log_2_ gene expression (CPM) of platelet-specific genes (*PF4*, *PPBP* and *NRGN*) (Table S3) across clusters of blood immune cells. In *(**C**), (**D**), (**E**)* and *(**F**)* P values were calculated with Wilcoxon test, taking Neu 5 cluster as the population of reference for each pairwise comparison with other clusters. P values were adjusted with Bonferroni test. Adjusted p-values (Adj P) were displayed on graphs, **** Adj P ≤ 0.0001, *** Adj P ≤ 0.001, ** Adj P ≤ 0.01, * Adj P ≤ 0.05, ns = Adj P > 0.05.

To validate that this Neu 5 cluster of platelet gene-expressing neutrophils containing NPAs was not exclusive to this particular scRNAseq dataset, we next investigated for the presence of platelet gene-expressing neutrophils in a public scRNAseq dataset of HDs’ whole blood leukocytes (GSE145230). By performing the same analysis as for the NSCLC patients scRNAseq dataset (GSE127465), we were able to identify a cluster of neutrophils (Neu 17) located in between the platelet cluster and the remaining neutrophil clusters (Fig. S1A-B). Consistent with what we previously found in the NSCLC patients scRNAseq dataset, a fraction of Neu 17 cells was expressing high level of neutrophil (Fig. S1C-E) and platelet-specific genes (Fig. S1D-F).

In order to rule out that upregulation of platelet genes in neutrophils is not a consequence of neutrophil activation, we next tested whether the enrichment of the platelet-specific genes signature “Raghavachari” was increased upon neutrophil activation. We used public transcriptomic data of *in vitro* activated neutrophils by either GM-CSF [45] or plasma from septic patients [46] and we show that the enrichment score of the platelet signature in neutrophils did not significantly differ upon activation as compared to controls (Fig. S2). The expression of platelet genes in Neu 5 (in GSE127465) or Neu 17 (GSE145230) clusters seems therefore not to result from neutrophil activation. Collectively, all these results strongly suggest that Neu 5 and or Neu 17 neutrophil cluster displaying a high expression of platelet genes most likely contained NPAs.

### Gene signature of neutrophils from NPAs reveals enhanced degranulation, chemotaxis and trans-endothelial migration capacities

To go deeper into the characterization of NPAs from NSCLC patients’ whole blood leukocytes at the mRNA level, we focused our analysis on NSCLC patients’ scRNAseq dataset (GSE127465). We noticed that only a fraction of cells of the Neu 5 cluster expressed platelet specific genes (*PF4*, *PPBP* and *NRGN*) (Fig. 3F). When we re-projected cells for each individual neutrophil cluster, keeping the same UMAP parameters used to discriminate each individual subset of leukocytes, we could show that the Neu 5 cluster was more heterogeneous than the other neutrophil clusters based on visual cell dispersion (Fig. 4A and Fig. S3). We selected the 48 cells co-expressing the 3 platelet specific genes (*PF4*, *PPBP* and *NRGN*) at a level above 1 log_2_ CPM (Fig. 4A) (named “Neu 5 PPBP^high^ PF4^high^ NRGN^high^”) out of the 437 of the whole Neu 5 cluster. 48 cells out of the 8873 neutrophils analyzed represents 0,54% which is consistent with the median frequency of NPAs among all neutrophils (0.85%) found in our cohort of NSCLC patients (Fig. 1D and Table S1).

**Figure 4.**
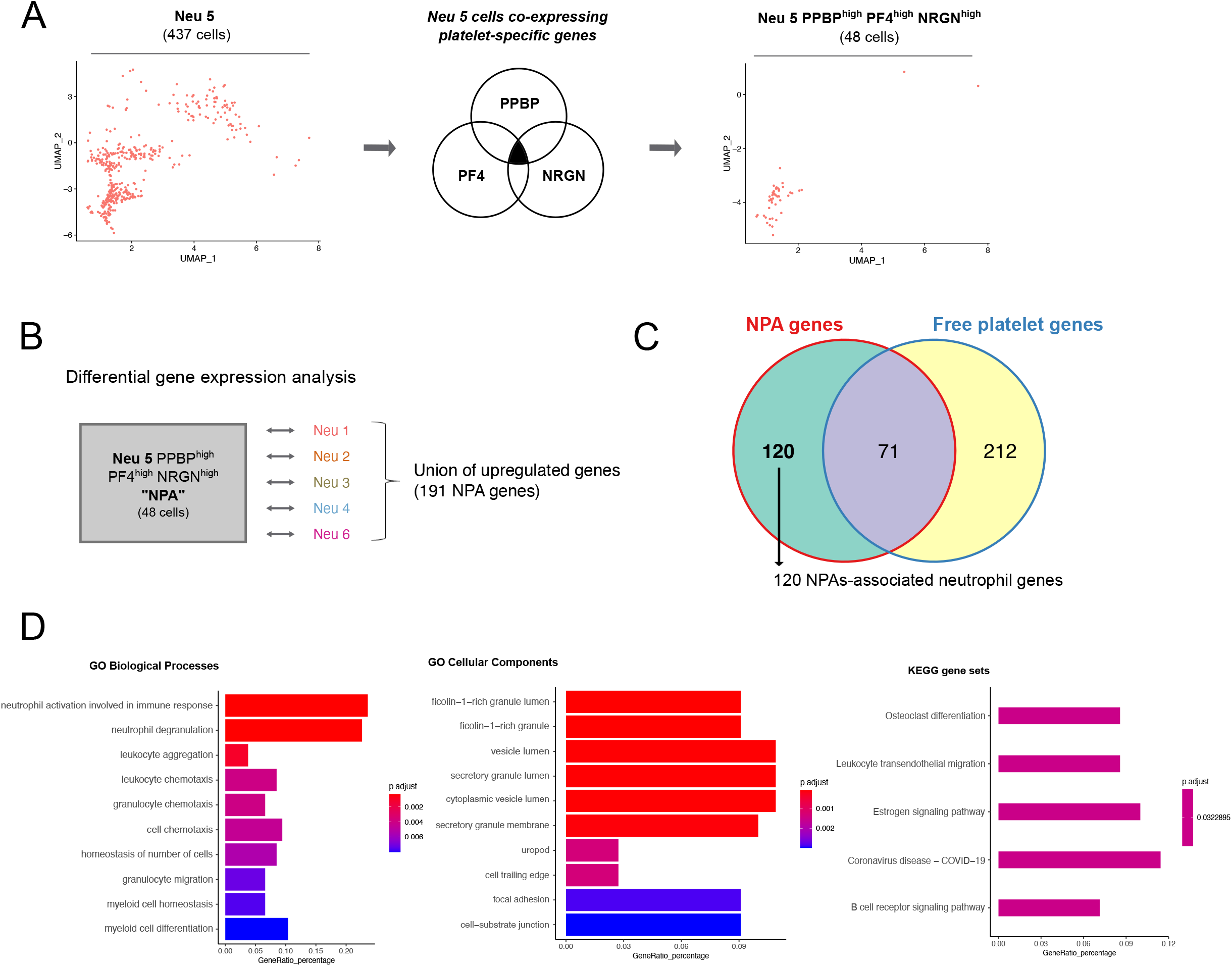
NPA gene signature is enriched for neutrophil degranulation, chemotaxis and trans-endothelial migration GO terms, as opposed to the other neutrophil subsets. *(**A***) Strategy to make the Neu 5 population more homogeneous (public scRNAseq data from NSCLC patients’ whole blood leukocytes - GSE127465). Left panel represents the two dimensional-UMAP projection plot of cells from Neu 5 cluster (comprising 437 cells), based on the UMAP parameters used to discriminate the major subsets of leukocytes. Middle panel shows the selection of Neu 5 cells based on the co-expressed high level of the three platelet specific genes (*PPBP, PF4* and *NRGN*) above 1 log_2_ gene expression (CPM). Right panel represents the two dimensional-UMAP projection plot of Neu 5 cells co-expressing *PPBP, PF4* and *NRGN* genes (comprising 48 cells annotated Neu 5 PPBP^high^ PF4^high^ and NRGN^high^). Cells were plotted based on the UMAP parameters used to discriminate the major subsets of leukocytes. *(**B**)* Differential gene expression approach to get the union of upregulated genes in Neu 5 PPBP^high^ PF4^high^ NRGN^high^ (annotated NPA cluster) as opposed to each of the remaining neutrophil clusters (Neu 1, Neu 2, Neu 3, Neu 4 and Neu 6). The statistical test used in the differential gene expression analysis was the Wilcoxon rank-sum (Mann–Whitney) test. *(**C**)* Venn diagram representing mutually and exclusive upregulated genes between the NPA cluster (union of upregulated genes in NPA cluster compared to each of the remaining neutrophil clusters – see Table S4) and the platelet cluster (Table S3). *(**D**)* Top 10 most-enriched GO Biological Processes, Cellular Components and KEGG gene sets calculated based on the 120 specific upregulated genes in NPA compared to platelet cluster (Table S5).

To next determine if neutrophils from the NPA cluster were distinct from any other subset of neutrophils, we first performed a differential gene expression analysis of the NPA cluster (Neu 5 PPBP^high^ PF4^high^ NRGN^high^ cells) in comparison to each individual neutrophil clusters (Fig. 4B). We then took the union of upregulated genes in NPA cluster from each pairwise comparison, therefore yielding 191 genes. We then removed the platelet genes (Fig. 4C) and identified 120 non-platelet genes discriminating NPAs-derived neutrophils from any other subset of neutrophils.

We next performed a pathway analysis and found a significant enrichment of neutrophil degranulation/chemotaxis-related GO biological process, secretory granules-related GO cellular component and trans-endothelial migration-related KEGG gene set (Fig. 4D). Collectively, these data demonstrate that NPAs-derived neutrophils display enhanced degranulation, chemotaxis and trans-endothelial migration functionalities.

### Specific NPAs-associated neutrophil signature is associated with a worse prognosis in pancreatic adenocarcinoma and liver hepatocellular carcinoma patients

Next we addressed the prognostic significance of neutrophils from NPAs across the different tumor types available in The Cancer Genome Atlas (TCGA). The first step was to generate a specific NPAs-associated neutrophil signature excluding any other genes expressed in the other non-neutrophil blood immune cell type and platelets (Fig. 5A) yielding 16 non-platelet NPAs-associated neutrophils specific genes (Fig. 5B). Using scRNAseq data from human NSCLC tumor samples (discovery dataset) matching previously-used patients’ whole bloods [31], we then eliminated *RBPJ*, *IDH2*, *HELLPAR* and *CCNI* that were highly expressed by multiple other tumor-associated cell types (Fig. S4), thus yielding a final 12-genes NPAs-associated neutrophil signature (Fig. 5B). We next validated the specificity of this new NPAs-associated neutrophil signature in the discovery dataset (scRNAseq dataset from NSCLC tumors) by showing that cells coexpressing at least 3 out of the 12-genes NPA-associated neutrophil signature were significantly more enriched in tumor-associated neutrophils than any other major cell types present in the tumor microenvironment (Fig. 5C).

**Figure 5.**
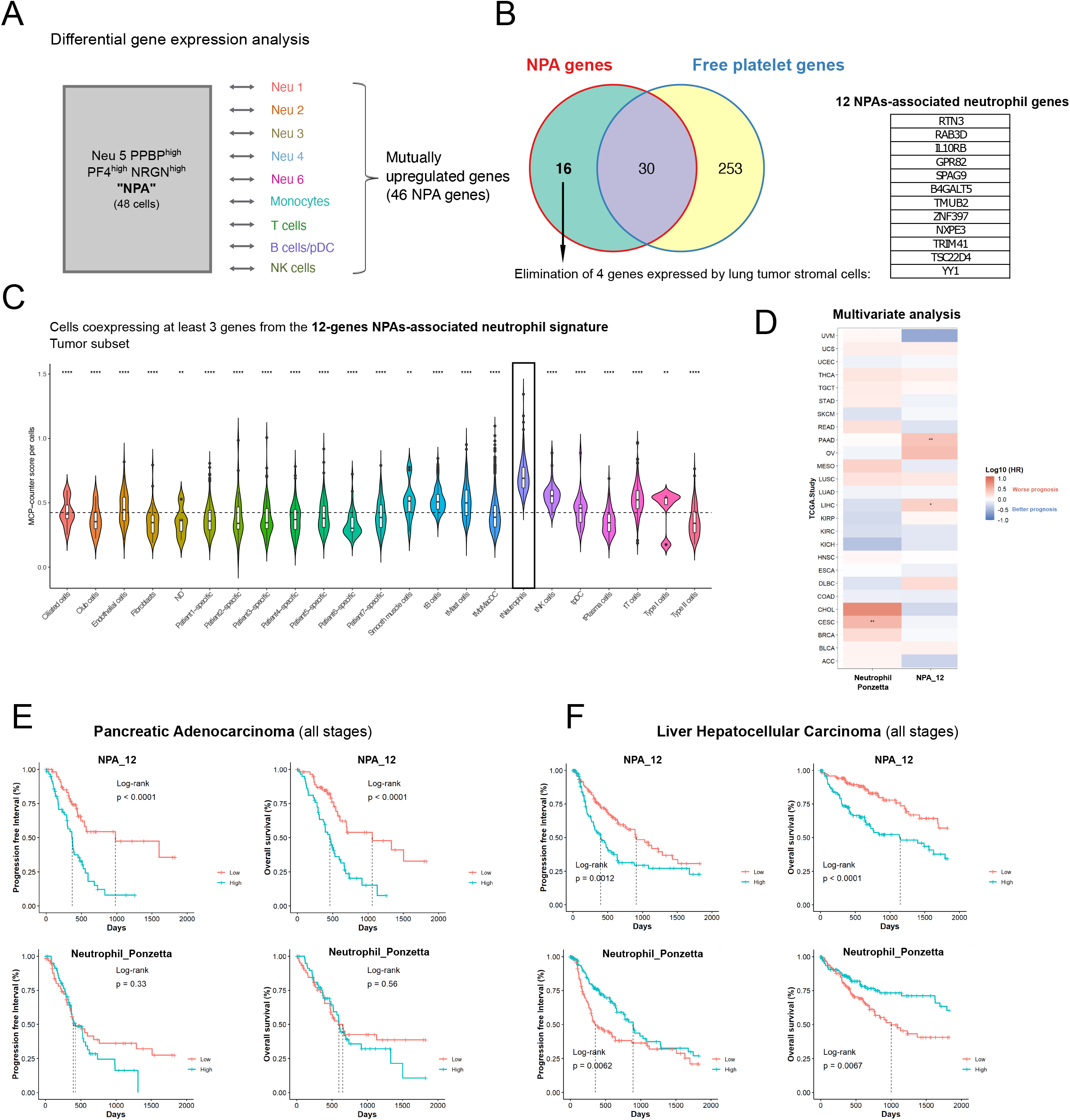
Specific NPA gene signature is associated with a worse prognosis in pancreatic adenocarcinoma and liver hepatocellular carcinoma patients. *(**A**)* Differential gene expression approach was used to get the list of mutually upregulated genes in Neu 5 PPBP^high^ PF4^high^ NRGN^high^ (annotated NPA cluster) across all pairwise comparison with the remaining neutrophil clusters (Neu 1, Neu 2, Neu 3, Neu 4 and Neu 6) and non-neutrophil clusters (Monocytes, T cells, B cells/pDC and NK cells) based on public scRNAseq data from NSCLC patients’ whole blood leukocytes - GSE127465. The statistical test used in the differential gene expression analysis was the Wilcoxon rank-sum (Mann–Whitney) test. *(**B**)* Venn diagram representing mutually and exclusive upregulated genes between the NPA cluster (mutually upregulated genes in NPA cluster across all pairwise comparison with the remaining neutrophil clusters and non-neutrophil clusters – see Table S6) and the platelet cluster (Table S3), yielding 16 upregulated genes specific to NPAs as compared to free platelet cluster. 4 genes out of the 16, were removed as we they were highly expressed by lung tumor stromal cells (based on public scRNAseq data from human NSCLC tumors (GSE127465)), giving rise to the 12 gene NPA signature (listed in a table). *(**C**)* Violin plot representing the MCP counter score per cell for each stromal non-immune cell types (Ciliated cells, Club cells, Endothelial cells, Fibroblasts, Smooth muscle cells, Type I pneumocytes and Type II pneumocytes), stromal immune cell types (B cells, Mast cells, Monocytes/Macrophages/DC (Dendritic Cells), Neutrophils, NK (Natural Killer) cells, pDC (Plasmacytoid Dendritic cells), Plasma cells and T cells) and patient-specific tumor cell clusters, based on public scRNAseq data from human NSCLC tumors (GSE127465). All cell types were labeled based on author’s annotations (GSE127465). *(**D**)* Heatmap summarizing results of the multivariate Cox analysis of prognostic values of various immune signatures across all available TCGA cohorts. Y axis correspond to TCGA cohorts. Each line corresponds to a particular cancer type annotated with the TCGA short name. X axis corresponds to the canonical neutrophil signature (annotated “Neutrophil_Ponzetta” – see Table S2) and the 12-genes NPAs-associated neutrophil signature (annotated “NPA_12” – see Figure 5.B). HR corresponds to the hazard ratio expressed in log_10_. Positive HR values (in red) correspond to a worse prognosis and negative HR values (in blue) correspond to a better prognosis. * refers to an adjusted p-value < 0.05. ** refers to an adjusted p-value < 0.01. *(**E-F**)* Univariate survival analysis in pancreatic adenocarcinoma (PAAD) and Liver Hepatocellular Carcinoma (LIHC) cancer patients (all stages) using Kaplan Meier method. Cohort of PAAD and LIHC cancer patients were cut at tercile values based on mean expression of “Neutrophil_Ponzetta” or “NPA_12” signature, yielding two groups (highest tercile or lowest tercile for expression of the signature of interest).

We next evaluated the prognostic value of the 12-genes NPAs-associated neutrophil signature (Fig. 5B) and the canonical neutrophil signature “Ponzetta” (Table S2) with a pan-cancer approach in the TCGA database. By performing a multivariate Cox analysis, we did not find any association between the NPAs-associated neutrophil signature and the prognosis of either types of NSCLC cancer patients, lung squamous cell carcinoma (LUSC) and lung adenocarcinoma (LUAD) (Fig. 5D). Nevertheless, we could find that the signature was statistically associated with a worse progression-free interval (PFI) (p-value < 0.001, Hazard Ratio (HR) = 2.9) and overall survival (OS) (p-value < 0.001, HR = 2.6) in pancreatic adenocarcinoma cancer patients (PAAD) (Fig. 5D and Fig. S5A), as well as in liver hepatocellular carcinoma cancer patients (LIHC) (PFI: p-value = 0.063, HR = 1.45; OS: p-value = 0.004, HR = 2.1) (Fig. 5D and Fig. S5B). The canonical neutrophil signature “Ponzetta” could not recapitulate such significant prognostic values in PAAD and LIHC reinforcing the idea that NPAs-associated neutrophils is a unique population of neutrophils with different outcomes (Fig. 5D). Kaplan Meier univariate analysis confirmed the poorer prognostic values of the 12-genes NPAs-associated neutrophil signature in PAAD (Fig. 5E) and LIHC cancer patients (Fig. 5F). Whereas the canonical neutrophil signature “Ponzetta” did not show statistically significant association with prognosis in PAAD cancer patients (Fig. 5E), it was associated with an opposite outcome in LIHC cancer patients (Fig. 5F). This implies again that NPAs-associated neutrophils are distinct from canonical neutrophils. Collectively, our results therefore suggest that a subset of neutrophils, corresponding to NPAs-associated neutrophils, when present in tumors may be detrimental in PAAD and LIHC cancer patients.

## Discussion

To our knowledge, our study is the first to report the identification and characterization of NPAs at the transcriptomic level, based on scRNAseq data from human whole blood leukocytes. Based on their transcriptomic profile, we found that neutrophils bound to NPAs displayed enhanced degranulation, chemotaxis and trans-endothelial migration features. Our results collectively showed that neutrophils from NPAs represent a distinct subset of neutrophils, related to LDNs. Lastly, using TCGA database, we showed that our specific NPAs-associated neutrophil signature was associated with a worse prognosis in PAAD and LIHC cancer patients. Hence, NPAs may hold clinical utility as novel non-invasive blood prognostic biomarker in cancer patients with solid tumors.

Using ISX, we were able to distinguish and quantify accurately NPAs in our cohort of NSCLC cancer patients and HDs. Although we did not find a significant difference in the percentage of NPAs between NSCLC cancer patients and HD, one NSCLC patient displayed more than a three-fold increase of NPA proportion compared to the mean frequency of NPAs in NSCLC patients and HDs (Fig. 1D). Interestingly, this patient also displayed the highest number of different metastatic sites (Table S1), as well as an extremely high NLR value of 34.56 (Table S1). This is consistent with the poorer clinical outcome of patients with elevated NLR values, including NSCLC patients [47]. This high NLR value was mostly explained by a massive increase in the absolute neutrophil count (ANC), rather than a decrease in the absolute lymphocyte count. Mounting evidence have shown that high ANC alone, was also significantly associated with poorer prognosis in NSCLC cancer patients [48,49], including in response to immune checkpoint inhibitors (ICIs) [50–52]. It would be of great interest in future studies to determine if the percentage of circulating NPAs, in comparison to the other cost-effective and non-invasive NLR and ANC, is more accurately associated with response to immunotherapy.

By performing a Ficoll density gradient, we showed that NPAs define a subset of LDNs but are absent in NDN fraction, illustrating that platelets preferentially bind to LDN in contrast to NDN (Fig. 3B-C-D). Several studies have reported a link between LDNs and cancer progression [47]. Many research groups indeed reported that LDNs displayed enhanced T-cell suppressive properties, expanded in cancer patients’ blood as opposed to healthy donors [11–14,53]. Recent evidence have shown that high LDN expansion in cancer patients’ blood was associated with worse prognosis [11,12], including NSCLC patients [12]. Nevertheless, it remains to be known why platelets bind preferentially to LDNs. The lower density of neutrophil bound to platelet may be a consequence of platelet binding. Since NDN activation is sufficient to convert NDNs into LDNs [41], one can hypothesize that platelets interacting with NDN activate and convert them into LDNs. It is also likely that platelets may behave as floats changing NDNs’ density and retaining them at the surface of the Ficoll gradient, explaining their recovery within peripheral blood mononuclear cells. Although we did not document a significant increase of NPAs frequency in patient, likely as a consequence of the small size of the cohort, the fact that LDNs increase in cancer patients strongly suggests that the frequency of NPAs also increases in cancer patients.

Although unexpected, this study identified a yet unreported new subset of neutrophils in scRNAseq data with high platelet gene expression that most likely correspond to NPAs. We ruled out the possibility that platelet genes could be expressed by a subset of activated neutrophils as we did not find enrichment of platelet signature in stimulated neutrophils (Fig. S2). Moreover, it is also unlikely that neutrophils expressing platelet genes correspond to a common progenitor of megakaryocyte (platelet-producing cell) and neutrophils, as the closest common progenitor between the two is the common myeloid progenitor (CMP), which gives rise to all myeloid cells [54]. High expression of neutrophil-specific genes at this stage of differentiation is therefore unlikely to occur. Taken together, platelet genes are more likely to be expressed exogenously by neutrophils through aggregated-platelets. It is however plausible that platelets could have been isolated in the same droplet than neutrophils, without any interaction. The potential random contamination of platelets during single cell isolation of immune cells would have yielded cluster of cells expressing platelet-specific genes in all immune cell types. However, expression of platelet specific genes was mostly observed in neutrophils as opposed to other major immune cell types (Fig. 3F). To note that some monocytes also displayed expression of platelet specific genes which is consistent with the fact that platelets may also aggregate with circulating monocytes [18]. Interestingly, we found that the frequency of neutrophil co-expressing the 3 platelet specific genes (*PF4*, *PPBP* and *NRGN*) was very similar to the one we accurately determined by ISX in our cohort of NSCLC patients. This further supports that the cluster of neutrophils with high expression of platelet genes identified in Zilionis’ scRNAseq dataset (GSE127465) corresponds to NPAs.

In contrast to the other subsets of circulating neutrophils, we found that neutrophils from NPAs displayed enhanced degranulation, chemotaxis and trans-endothelial migration features, based on the specific transcriptomic signature of NPAs. This is consistent with previous reports showing that platelet binding to neutrophils boosted their potential to release granules [22], increased their capacity to transmigrate through endothelial cells [25,26] to exacerbate inflammation thereby promoting disease progression [21,27–29]. One can speculate that if neutrophils from NPAs display higher transmigration potential, they would infiltrate tumors more efficiently and then fuel tumor growth and metastasis.

We found that NPA signature was highly detected in few tumor types in the TCGA data base and was strongly associated with a poorer prognosis in PAAD and LIHC cancer patients. This is consistent with the tumor-promoting role of TAN documented in pancreatic [55,56] and liver [57] cancer. Interestingly, we found that the canonical neutrophil signature “Ponzetta” could not recapitulate such significant poorer prognostic values in PAAD and LIHC, reinforcing the idea that neutrophil from NPA would account for most of tumor-promoting role of TAN in such cancer types. Moreover, we provided evidence that NPAs-associated neutrophils more likely resemble LDNs (Fig. 2) that are described to possess tumor-promoting functions [11,13], supporting the idea that NPAs-associated neutrophils once becoming TANs after tumor infiltration would fuel tumor growth.

Beyond their potential role in cancer, recent studies have reported an increase in the percentage of circulating NPAs in COVID-19 patients in contrast to HDs [58,59]. Interestingly, patients who had severe forms of COVID-19 displayed a significant higher level of NPAs in contrast to patients who had moderate forms [59]. We showed that NPAs displayed enhanced degranulation/activation, based on their specific transcriptomic signature (Fig. 4D). This is in accordance with a recent report showing in bulk granulocyte RNAseq data from COVID-19 patients an increase expression of granulocyte activation-associated factors in severe COVID-19 patients in contrast to mild ones [60]. Whether NPAs may not just be a consequence but also a cause of COVID-19 severity remains to be investigated in future studies.

## Declarations

### Ethics approval and consent to participate

Metastatic NSCLC patients (Table S1) included in the LIBIL clinical trial (NCT02511288) at the Léon Bérard Hospital as well as healthy donors from Etablissement Français du Sang provided informed consent regarding the use of their blood for research purpose.

### Consent for publication

Not applicable

### Availability of data and materials

The publicly available datasets analyzed during the current study are available from the *GEO repository*

GSE15139 (https://www.ncbi.nlm.nih.gov/geo/query/acc.cgi?acc=GSE15139)

GSE49757 (https://www.ncbi.nlm.nih.gov/geo/query/acc.cgi?acc=GSE49757)

GSE79404 (https://www.ncbi.nlm.nih.gov/geo/query/acc.cgi?acc=GSE79404)

GSE127465 (https://www.ncbi.nlm.nih.gov/geo/query/acc.cgi?acc=GSE127465)

GSE145230 (https://www.ncbi.nlm.nih.gov/geo/query/acc.cgi?acc=GSE145230)

### Authors’ contributions

Conceptualization: PL, CC and M-CM; methodology: PL, MA, SD, VA, LM, MH, HH-V, AV, CC and M-CM; formal analysis: PL, MA, SD, VA, LM and MH; investigation: PL, MA, SD, VA, LM, MH, AS, HH-V, AV, CC and M-CM; resources: AS, CC and M-CM; writing—original draft: PL; writing—review and editing: MA, SD, VA, LM, CC and M-CM; visualization: PL, MA, SD, VA, LM and MH; supervision: CC and M-CM; funding acquisition: PL, AV, CC and M-CM. All authors read and approved the final manuscript.

## Acknowledgements

We thank Laurie Tonon, Janice Kielbassa, Manuela Pereira-Abrantes, Jenny Valladeau-Guilemond, Roxane Pommier and Elisa Gobbini for careful reading of the manuscript, helpful comments and suggestions. We thank all members of the Caux team for their help and critical scientific comments. We want to thank the staff of the core facilities at the Cancer Research Center of Lyon (CRCL) for technical assistance, especially the cytometry core facility, the biological resource center (BRC) of the Centre Léon Bérard (CLB) for providing human samples, and the cytology lab (Emilie Josserand, Amandine Boyer) for helping us in performing May-Grünwald-Giemsa staining. We also thank Sarah Barrin and Justine Berthet for helping us taking images.

## Supplemental Figures

**Figure S1. Identification of a cluster of neutrophils expressing platelet genes in public scRNAseq data from healthy donors’ peripheral blood leukocytes.**

*(**A-B**)* Two dimensional-UMAP representation of re-clustered NSCLC patients’ whole blood leukocytes from scRNAseq data (GSE145230). *(**A**)* Major immune cell types were labeled using Clustifyr and SingleR annotation tools from pre-selected immune cell signatures. *(**B**)* Each of the 22 clusters defined by the analysis were labeled based on the type of immune cell. The 6 clusters of neutrophils were annotated as follow: Neu 0, Neu1, Neu 6, Neu 15, Neu 17 and Neu 18. Dark arrows show Neu 17 neutrophil cluster of interest displaying high expression of platelet genes. *(**C**)* Violin plot representing ssGSEA enrichment score per cell of the Ponzetta neutrophil signature (Table S2) across all clusters of blood immune cells. *(**D**)* Violin plot representing ssGSEA enrichment score per cell of the Raghavachari platelet signature (Table S2) across all clusters of blood immune cells. *(**E**)* Violin plots representing the log_2_ gene expression (count per million, CPM) of neutrophil-specific genes (*CXCR2*, *CSF3R* and *FCGR3B*) (Table S2) across all clusters of blood immune cells. *(**F**)* Violin plots representing the log_2_ gene expression (CPM) of platelet-specific genes (*PF4*, *PPBP* and *NRGN*) (Table S3) across clusters of blood immune cells.

In *(**C**), (**D**), (**E**)* and *(**F**)* P values were calculated with Wilcoxon test, taking Neu 5 cluster as the population of reference for each pairwise comparison with other clusters. P values were adjusted with Bonferroni test. Adjusted p-values (Adj P) were displayed on graphs, **** Adj P ≤ 0.0001, *** Adj P ≤ 0.001, ** Adj P ≤ 0.01.

**Figure S2. Enrichment analysis of platelet signatures in stimulated versus unstimulated neutrophils.**

*(**A**)* Box plot representing ssGSEA enrichment scores of two independent platelet signatures Raghavachari platelet signature (Table S2) in GM-CSF-treated neutrophils (“treated”) versus untreated neutrophils (“Control”). Blood from 3 HDs were used for each group (GSE15139). Differential enrichment of platelet signature between “treated” and “control” groups was assessed by the Wilcoxon rank-sum (Mann–Whitney) statistical test (p-value = 0.4). *(**B**)* Box plot representing ssGSEA enrichment scores of the Raghavachari platelet signature (Table S2) in neutrophils treated either with plasma from septic patients (“Septic plasma”, n = 35) or from HDs (“Uninfected plasma”, n=19) (GSE49757). Differential enrichment of platelet signature between “Septic plasma” and “Uninfected plasma” groups was assessed by the Wilcoxon rank-sum (Mann– Whitney) statistical test (p-value = 0.05202).

**Figure S3. Two dimensional-UMAP representation of each neutrophil cluster.**

Two dimensional-UMAP representation of individual neutrophil clusters (from NSCLC patients’ whole blood leukocytes from scRNAseq data (GSE127465)) whose projection was based on UMAP parameters used to discriminate the major subsets of leukocytes.

**Figure S4. Two dimensional-UMAP representation of NSCLC patients’ tumor scRNAseq data.** Expression of each of the 16 genes of the NPA signature (Table S5) plotted on the two dimensional-UMAP representation of NSCLC patients’ stromal and patient-specific tumors from scRNAseq data (GSE127465)).

**Figure S5. Multivariate analysis (Forest plot) of the prognostic impact of the 12-genes NPAs-associated neutrophil signature.**

Multivariate analysis taking into account age and stage as cofounding factors. Cohort of PAAD *(A)* and LIHC *(B)* cancer patients were cut at tercile values based on mean expression of “NPA_12” signature. For both cohorts, progression free-interval (panel on the left) and overall survival (panel on the right) were assessed.

## Supplemental Tables

**Table S1. Blood-related biological information and clinical features of NSCLC patients and healthy donors.**

**Table S2. Published platelet and neutrophil-specific signatures.**

**Table S3. List of upregulated genes in free platelet cluster as compared to all other leukocytes.**

Based on public scRNAseq data from NSCLC patients’ whole blood leukocytes - GSE127465.

**Table S4. List of upregulated genes in Neu 5 as compared to all other leukocytes.**

Based on public scRNAseq data from NSCLC patients’ whole blood leukocytes - GSE127465.

**Table S5. Platelet gene-free specific NPA signature as opposed to the remaining neutrophils clusters.**

List of 120 genes of the NPA signature generated based on public scRNAseq data from NSCLC patients’ whole blood leukocytes - GSE127465.

**Table S6. NPAs-associated neutrophil signature as opposed to the remaining neutrophils clusters and all the other leukocytes.**

List of 16 genes of the NPA signature generated based on the public scRNAseq data from NSCLC patients’ whole blood leukocytes - GSE127465.

## List of abbreviations

ANC: Absolute neutrophil count
CPM: Count per million
GO: Gene ontology
HDs: Healthy donors
HNC: Head and neck cancer
HR: Hazard Ratio
ISX: ImageStream®X
LDNs: Low-density neutrophils
LIHC: Hepatocellular carcinoma cancer patients
LUAD: Lung adenocarcinoma
LUSC: Lung squamous cell carcinoma
NDNs: Normal-density neutrophils
NETs: Neutrophil extracellular traps
NLR: Neutrophils-to-lymphocytes ratio
NPAs: Neutrophil-platelet aggregates
NSCLC: Non-small-cell lung cancer
OS: Overall survival
PAAD: Pancreatic adenocarcinoma cancer patients
PCA: Principal component analysis
PCs: Principal components
PFI: Progression-free interval
RBC: Red blood cells
RT: Room temperature
ssGSEA: Single sample Gene Set Enrichment Analysis
TANs: Tumor-associated neutrophils
TCGA: The Cancer Genome Atlas
UMAP: Uniform manifold approximation and projection

