## Supplementary materials for "Gene signature of circulating platelet-bound neutrophils is associated with poor prognosis in cancer patients"

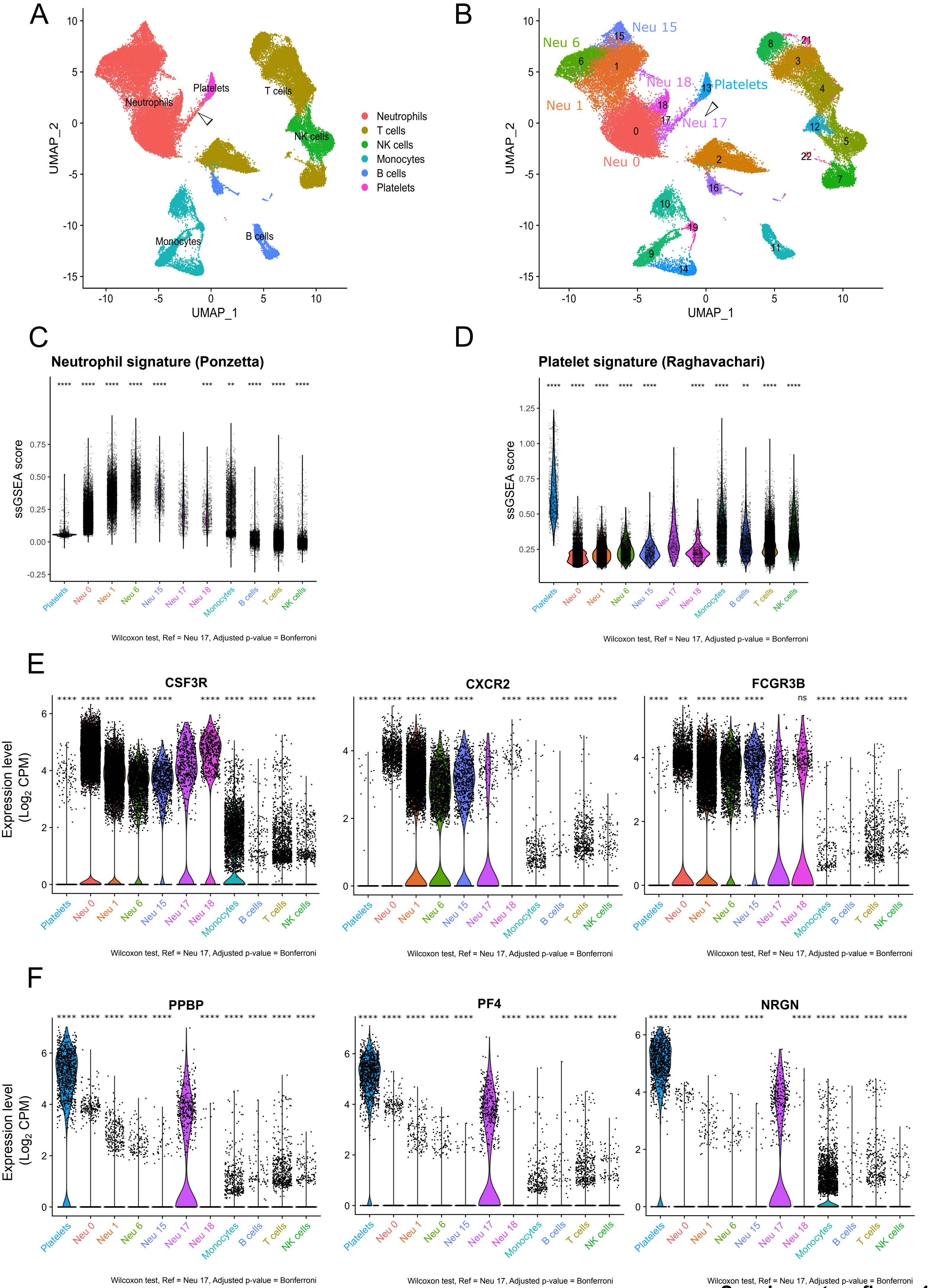

Supplementary figure 1

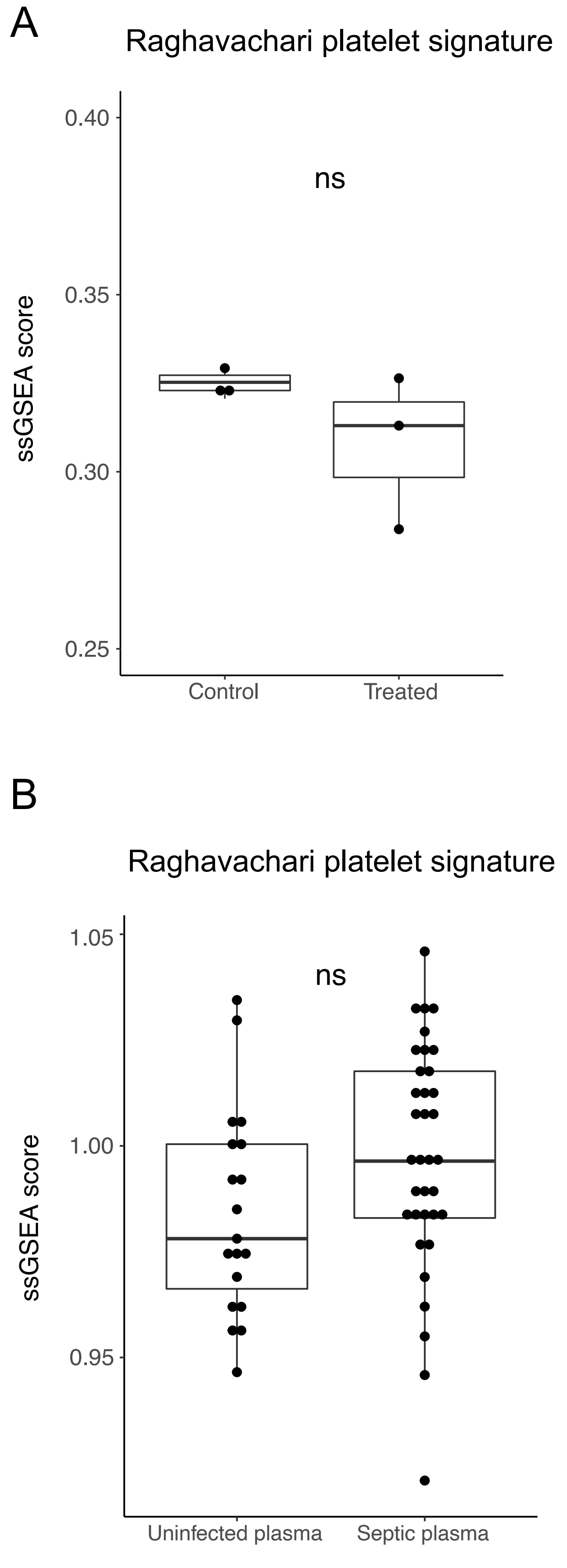

**Supplementary figure 2**

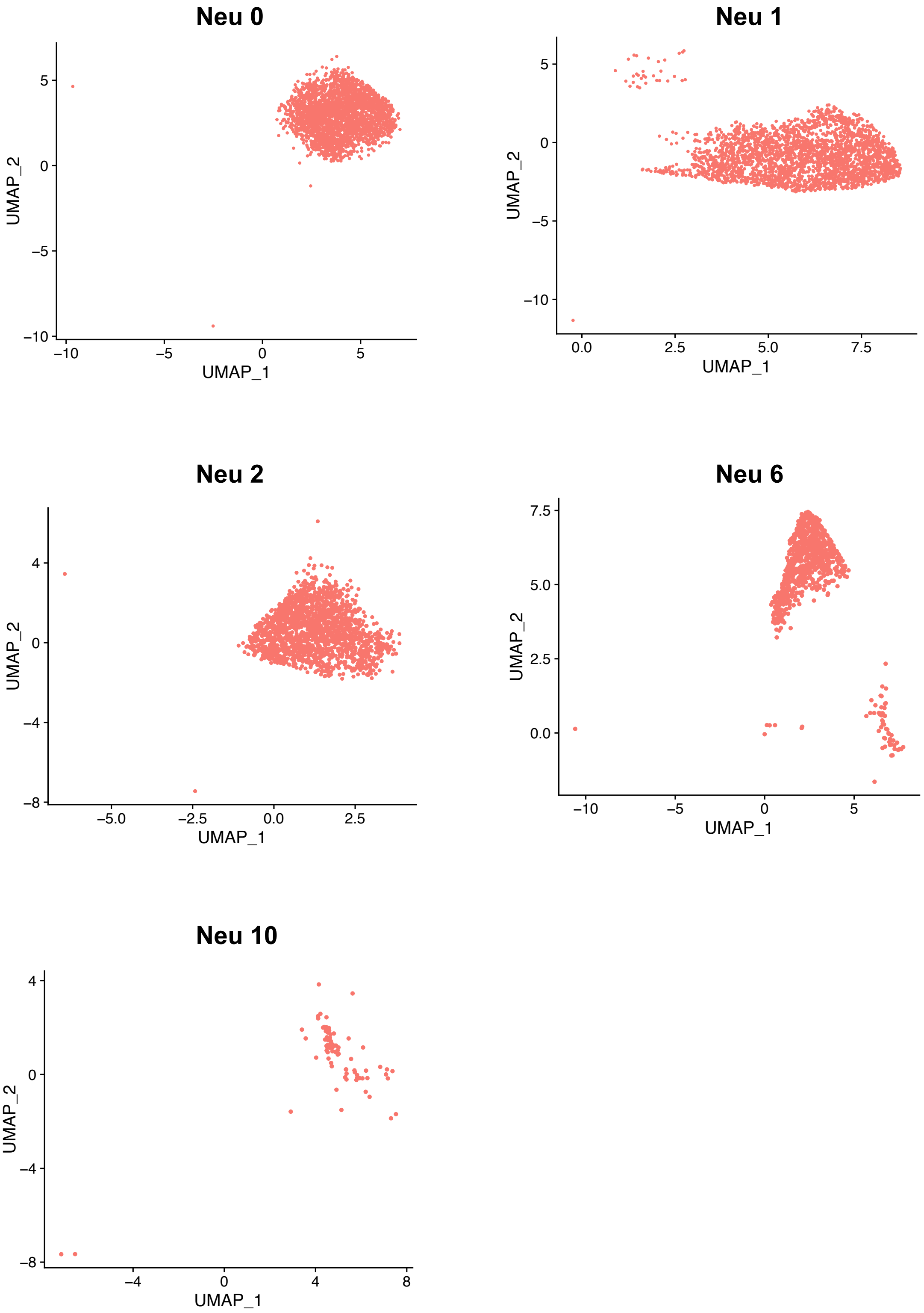

**Supplementary figure 3**

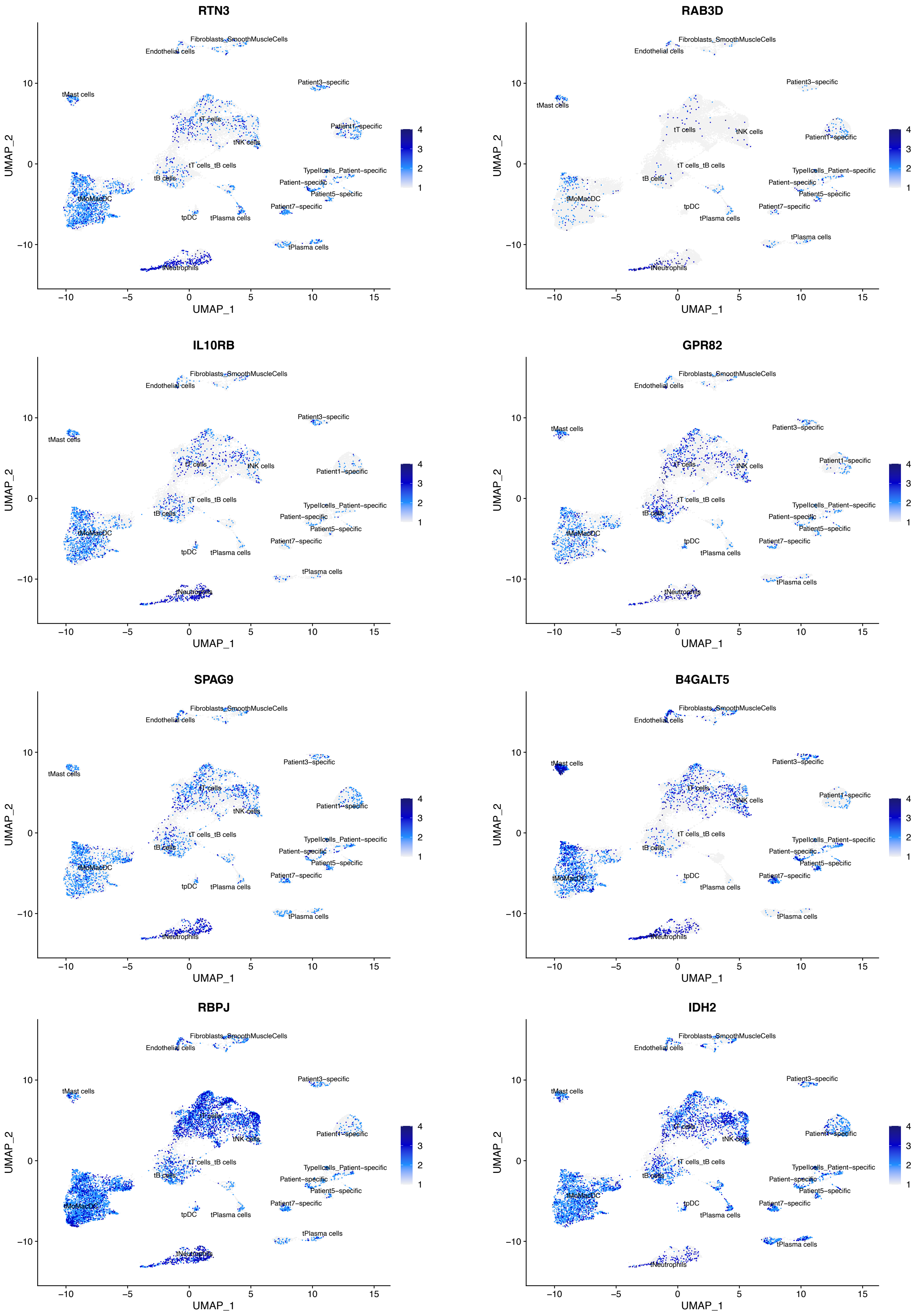

Supplementary figure 4

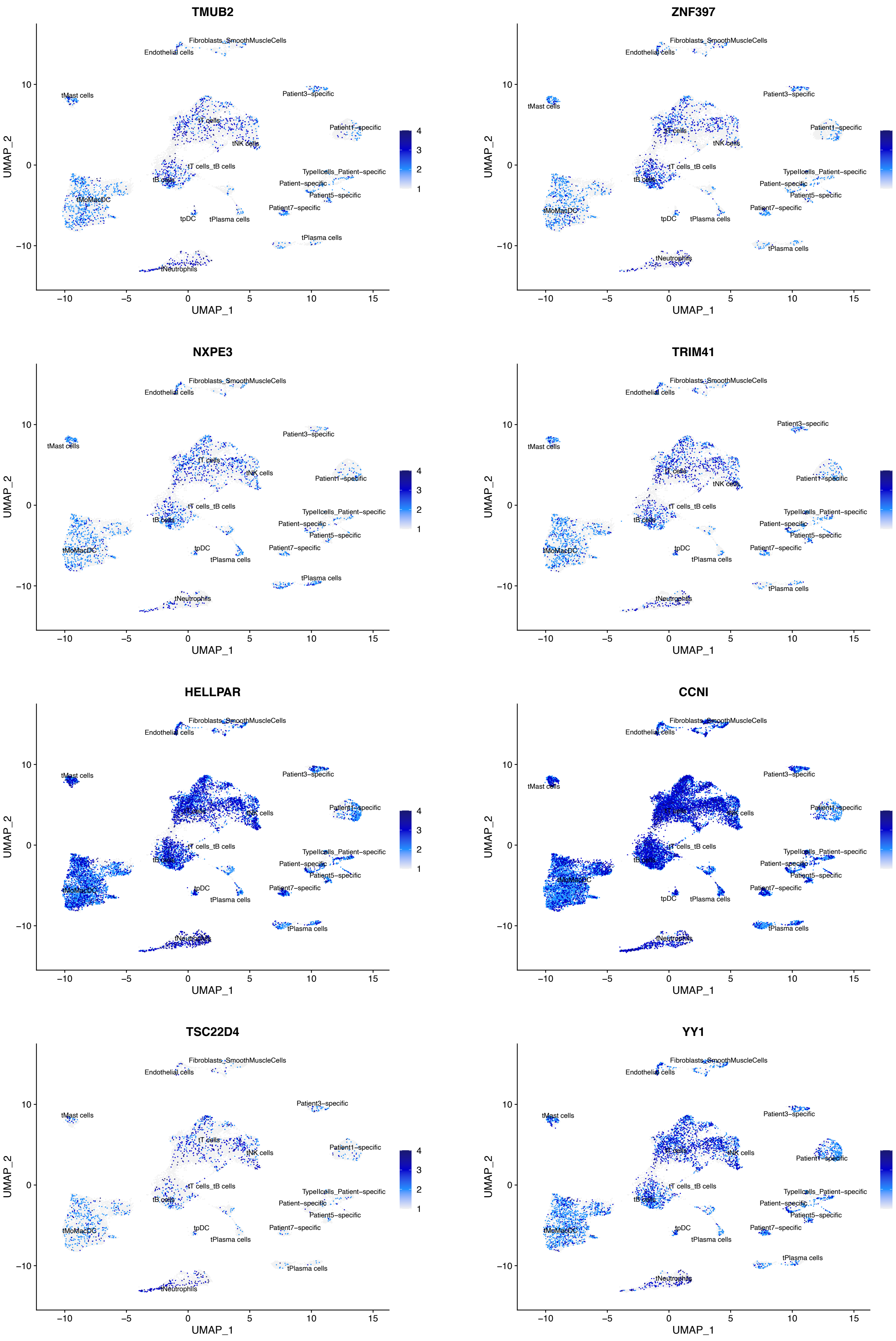

Supplementary figure 4 (next part)

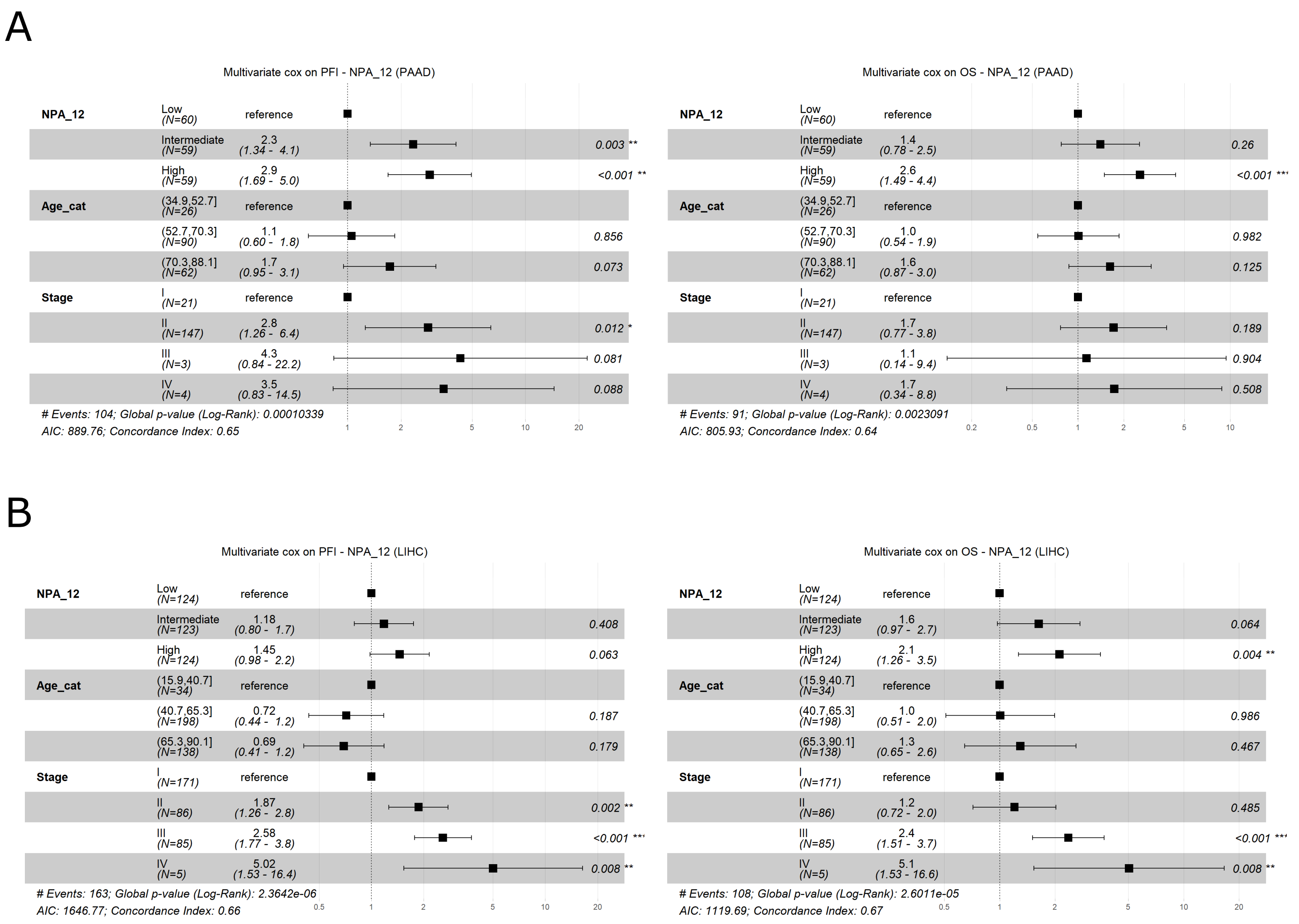

**Supplementary figure 5**

**Table S1. Clinical features of NSCLC patients and healthy donors.**

| Age at diagnosis (years) | Age interval (years) | Sex | Histology | Percentage of NPAs among neutrophils | Platelet number ($10^3/mm^3$) | Granulocytes number ($10^3/mm^3$) | Lymphocytes number ($10^3/mm^3$) | NLR (Gra/Lym) | Anti-platelet treatments | PD-L1 expression | Mutations | Metastatic site at diagnosis |
|---|---|---|---|---|---|---|---|---|---|---|---|---|
| 73 | >60 | Male | Lung adenocarcinoma | 0.8 | 200 | 4.1 | 0.8 | 5.1 | RESITUNE 75 | 100% | KRAS G12V | Lung, Lymph node |
| 54 | 51-60 | Male | Lung adenocarcinoma | 0.6 | 388 | 7.6 | 0.4 | 19.0 | none | 90% | No mutation | Lymph node, Adrenal Gland |
| 50 | 41-50 | Male | Lung adenocarcinoma | 0.7 | 229 | 6.0 | 2.9 | 2.1 | none | 100% | BRAF G469L | Lung, Lymph node, Bone |
| 49 | 41-50 | Female | Lung adenocarcinoma | 0.9 | 302 | 9.4 | 1.7 | 5.5 | none | 50% | Unknown | Lung, Lymph node, Pulmonary pleura, Brain |
| 55 | 51-60 | Male | Lung adenocarcinoma | 4.6 | 352 | 55.3 | 1.6 | 34.6 | none | 20% | KRAS Q61H | Liver, Bilateral Adrenal, Bone, Brain, Muscle, Mediastinal and Supraclavicular |
| 45 | 41-50 | Female | Lung adenocarcinoma | 1.4 | 257 | 2.6 | 0.6 | 4.3 | none | 20% | No mutation | Bone, Pleural |
| NA | 30- 40 | Female | Healthy donor | 0.8 | 260 | 3.3 | 2 | 1.7 | NA | NA | NA | NA |
| NA | 41-50 | Female | Healthy donor | 2.1 | 212 | 4.4 | 1.7 | 2.6 | NA | NA | NA | NA |
| NA | 41-50 | Female | Healthy donor | 1.6 | 256 | 5.6 | 2.6 | 2.2 | NA | NA | NA | NA |
| NA | 51-60 | Female | Healthy donor | 1.0 | 271 | 4.7 | 1.5 | 3.1 | NA | NA | NA | NA |
| NA | 51-60 | Female | Healthy donor | 0.4 | 236 | 3.9 | 1.5 | 2.6 | NA | NA | NA | NA |
| NA | 30- 40 | Male | Healthy donor | 1.2 | 133 | 3.9 | 2.1 | 1.9 | NA | NA | NA | NA |
| NA | 41-50 | Male | Healthy donor | 1.2 | 183 | 1.8 | 1.5 | 1.2 | NA | NA | NA | NA |
| NA | 41-50 | Male | Healthy donor | 1.1 | 169 | 2.7 | 1.2 | 2.3 | NA | NA | NA | NA |
| NA | 51-60 | Male | Healthy donor | 1.5 | 161 | 2.7 | 1.2 | 2.3 | NA | NA | NA | NA |
| NA | 51-60 | Male | Healthy donor | 0.7 | 232 | 3.1 | 1.8 | 1.7 | NA | NA | NA | NA |

### Table S2. Published platelet and neutrophil-specific signatures.

| Geneset name | RAGHAVACHARI_PLATELET_SIGNATURE | PONZETTA_NEUTROPHILS_SIGNATURE |
|---|---|---|
| PMID number | 17353439 | 31257026 |
| Description | abuntant and specific transcripts in blood | specific transcripts of blood neutrophils |
| Genes | APP | CSF3R |
| | ASAH1 | ALPL |
| | BEX3 | BST1 |
| | CCL5 | CD93 |
| | CTSA | CEACAM3 |
| | CTTN | CPPED1 |
| | CXCL5 | CREB5 |
| | DAPP1 | CRISPLD2 |
| | EIF2AK1 | CXCR1 |
| | ENDOD1 | CXCR2 |
| | F13A1 | CYP4F3 |
| | FHL1 | DYSF |
| | GLA | FCAR |
| | GNAS | FCGR3B |
| | GNG11 | FPR1 |
| | HBA1 | FPR2 |
| | HBB | G0S2 |
| | HBG1 | HIST1H2BC |
| | HIST1H2AC | HPSE |
| | HIST1H3H | KCNJ15 |
| | HIST2H2BE | LILRB2 |
| | ITGA2B | MGAM |
| | ITGB3 | MME |
| | ITGB5 | PDE4B |
| | KIF2A | S100A12 |
| | LEPROT | SIGLEC5 |
| | MAP3K7CL | SLC22A4 |
| | MAX | SLC25A37 |
| | MFAP3L | TECPR2 |
| | MLH3 | TNFRSF10C |
| | MMD | VNN3 |
| | MPP1 | |
| | MYLK | |
| | NAP1L1 | |
| | NRGN | |
| | ODC1 | |
| | PDLIM1 | |
| | PF4 | |
| | PF4V1 | |
| | PGRMC1 | |
| | PIP4K2A | |
| | PPBP | |
| | PPM1A | |
| | PRDX6 | |
| | PRKAR2B | |
| | PRKCB | |
| | PRUNE1 | |
| | PTGS1 | |
| | PTPN12 | |
| | RAB31 | |
| | RABGAP1L | |
| | RGS10 | |
| | RNF11 | |
| | RSU1 | |

| | |
|---|---|
| | RUFY1 |
| | RYBP |
| | SNCA |
| | SNN |
| | SPARC |
| | TAX1BP3 |
| | THBS1 |
| | TMEM140 |
| | TNFSF4 |
| | TPM1 |
| | TSC22D1 |
| | TUBA4A |
| | TUBB2A |
| | VCL |
| | WIPF1 |
| | YPEL5 |
| | ZNF185 |

**Table S3. List of upregulated genes in free platelet cluster as compared to all other leukocytes.**

| Genes | p_val | avg_logFC | pct.1 | pct.2 | p_val_adj |
|---|---|---|---|---|---|
| PF4 | 0 | 5.039797549 | 0.98 | 0.016 | 0 |
| PPBP | 0 | 4.837553685 | 0.949 | 0.018 | 0 |
| TUBB1 | 0 | 4.400580008 | 0.99 | 0.007 | 0 |
| NRGN | 0 | 4.287975287 | 0.99 | 0.045 | 0 |
| MPIG6B | 0 | 4.177363261 | 0.909 | 0.012 | 0 |
| F13A1 | 0 | 3.923542164 | 0.889 | 0.036 | 0 |
| ACRBP | 0 | 3.724207984 | 0.717 | 0.009 | 0 |
| CAVIN2 | 0 | 3.573199852 | 0.879 | 0.007 | 0 |
| SPARC | 0 | 3.447017281 | 0.859 | 0.01 | 0 |
| CLU | 0 | 3.332596351 | 0.768 | 0.027 | 0 |
| TSC22D1 | 0 | 3.214687883 | 0.646 | 0.016 | 0 |
| ITGA2B | 0 | 3.167009865 | 0.697 | 0.014 | 0 |
| PRKAR2B | 0 | 3.161044457 | 0.697 | 0.007 | 0 |
| GP9 | 0 | 3.139481586 | 0.727 | 0.002 | 0 |
| GRAP2 | 0 | 3.107960792 | 0.707 | 0.026 | 0 |
| MYL9 | 0 | 3.022337241 | 0.596 | 0.006 | 0 |
| SH3BGRL2 | 0 | 2.797828359 | 0.545 | 0.004 | 0 |
| CDKN1A | 0 | 2.795498714 | 0.485 | 0.013 | 0 |
| PDLIM1 | 0 | 2.732605884 | 0.485 | 0.012 | 0 |
| PTCRA | 0 | 2.709333321 | 0.556 | 0.001 | 0 |
| PTGS1 | 0 | 2.685495849 | 0.545 | 0.012 | 0 |
| CMTM5 | 0 | 2.665029949 | 0.505 | 0.003 | 0 |
| TMEM40 | 0 | 2.631153597 | 0.535 | 0.005 | 0 |
| TREML1 | 0 | 2.59846537 | 0.495 | 0.005 | 0 |
| SEPT5 | 0 | 2.569255869 | 0.646 | 0.013 | 0 |
| CTDSPL | 0 | 2.562497232 | 0.475 | 0.006 | 0 |
| MMD | 0 | 2.554421904 | 0.556 | 0.009 | 0 |
| BEX3 | 0 | 2.49623349 | 0.525 | 0.013 | 0 |
| HIST1H3H | 0 | 2.492951892 | 0.354 | 0.003 | 0 |
| C2orf88 | 0 | 2.461798158 | 0.475 | 0.004 | 0 |
| PF4V1 | 0 | 2.423733348 | 0.515 | 0.003 | 0 |
| ALOX12 | 0 | 2.372496933 | 0.343 | 0.001 | 0 |
| AL160400.1 | 0 | 2.356900047 | 0.374 | 0.006 | 0 |
| PDZK1IP1 | 0 | 2.320751836 | 0.404 | 0.003 | 0 |
| CD9 | 0 | 2.298487477 | 0.394 | 0.005 | 0 |
| SMOX | 0 | 2.274983246 | 0.424 | 0.002 | 0 |
| ITGB5 | 0 | 2.244073268 | 0.374 | 0.005 | 0 |
| ESAM | 0 | 2.181080132 | 0.343 | 0.003 | 0 |
| GNG11 | 0 | 2.129293415 | 0.414 | 0.004 | 0 |
| LTBP1 | 0 | 2.123161061 | 0.343 | 0.004 | 0 |
| GMPR | 0 | 2.109542511 | 0.354 | 0.004 | 0 |
| CTTN | 0 | 2.075451633 | 0.333 | 0.005 | 0 |
| AQP10 | 0 | 1.95266331 | 0.283 | 0.001 | 0 |
| GNAZ | 0 | 1.933152808 | 0.333 | 0.003 | 0 |
| SELP | 0 | 1.926906011 | 0.313 | 0.004 | 0 |
| TSPEAR-AS2 | 0 | 1.889751633 | 0.343 | 0.002 | 0 |
| PEAR1 | 0 | 1.888495643 | 0.303 | 0.003 | 0 |
| TSPAN33 | 0 | 1.856474821 | 0.333 | 0.003 | 0 |
| LGALS12 | 0 | 1.8276617 | 0.273 | 0.002 | 0 |
| AP001189.1 | 0 | 1.805543885 | 0.343 | 0.003 | 0 |
| ABLIM3 | 0 | 1.763964886 | 0.273 | 0.003 | 0 |
| HIST1H2BJ | 0 | 1.760156547 | 0.283 | 0.002 | 0 |
| CLDN5 | 0 | 1.751305664 | 0.263 | 0.003 | 0 |
| TUBA8 | 0 | 1.639131886 | 0.273 | 0.002 | 0 |
| MTURN | 2.07E-288 | 3.013839339 | 0.697 | 0.031 | 6.92E-284 |
| EHD3 | 1.04E-281 | 1.685946686 | 0.273 | 0.004 | 3.46E-277 |

| | | | | | |
|---|---|---|---|---|---|
| **TAL1** | 7.51E-274 | 1.708711418 | 0.253 | 0.003 | 2.51E-269 |
| **CLEC1B** | 5.45E-259 | 2.100710686 | 0.323 | 0.006 | 1.82E-254 |
| **MFAP3L** | 2.34E-250 | 1.615020293 | 0.283 | 0.005 | 7.82E-246 |
| **DAB2** | 1.34E-249 | 2.346604618 | 0.424 | 0.012 | 4.48E-245 |
| **ODC1** | 6.09E-234 | 2.486150623 | 0.525 | 0.021 | 2.03E-229 |
| **FHL1** | 7.86E-231 | 1.95601325 | 0.354 | 0.009 | 2.63E-226 |
| **MAP3K7CL** | 1.97E-222 | 2.640881295 | 0.485 | 0.019 | 6.58E-218 |
| **ELOVL7** | 1.14E-220 | 1.668605661 | 0.253 | 0.004 | 3.82E-216 |
| **GPX1** | 1.92E-212 | 3.386488363 | 0.909 | 0.083 | 6.43E-208 |
| **GAS2L1** | 2.62E-210 | 1.953750683 | 0.364 | 0.011 | 8.76E-206 |
| **PGRMC1** | 3.13E-208 | 2.309819183 | 0.465 | 0.019 | 1.04E-203 |
| **RUFY1** | 2.88E-202 | 2.782578644 | 0.636 | 0.038 | 9.63E-198 |
| **MARCH2** | 2.39E-200 | 2.456063388 | 0.606 | 0.035 | 7.98E-196 |
| **LMNA** | 4.46E-200 | 2.053870649 | 0.374 | 0.012 | 1.49E-195 |
| **SNCA** | 8.87E-197 | 1.875865346 | 0.293 | 0.007 | 2.97E-192 |
| **DMTN** | 4.81E-194 | 1.652369092 | 0.263 | 0.006 | 1.61E-189 |
| **RAB27B** | 8.80E-194 | 2.632716231 | 0.364 | 0.012 | 2.94E-189 |
| **ENDOD1** | 4.03E-168 | 2.02033365 | 0.333 | 0.011 | 1.35E-163 |
| **LGALSL** | 1.25E-156 | 1.888400647 | 0.283 | 0.009 | 4.17E-152 |
| **RGS10** | 2.42E-156 | 2.75998866 | 0.677 | 0.059 | 8.09E-152 |
| **NEXN** | 3.60E-155 | 2.173061083 | 0.293 | 0.01 | 1.20E-150 |
| **PCSK6** | 1.32E-150 | 1.517319887 | 0.263 | 0.008 | 4.42E-146 |
| **LINC00608** | 4.08E-148 | 2.141776737 | 0.414 | 0.021 | 1.36E-143 |
| **ZNF185** | 9.07E-146 | 2.074989811 | 0.384 | 0.018 | 3.03E-141 |
| **FAXDC2** | 9.27E-145 | 2.230206005 | 0.556 | 0.04 | 3.10E-140 |
| **ARHGAP18** | 3.36E-143 | 1.679119614 | 0.343 | 0.015 | 1.12E-138 |
| **VCL** | 1.88E-136 | 3.347463082 | 0.879 | 0.127 | 6.27E-132 |
| **YWHAH** | 3.81E-130 | 2.440954248 | 0.576 | 0.049 | 1.27E-125 |
| **SLA2** | 9.21E-126 | 2.176549419 | 0.384 | 0.021 | 3.08E-121 |
| **HIST1H2AC** | 4.76E-122 | 2.609776108 | 0.808 | 0.108 | 1.59E-117 |
| **THBS1** | 1.55E-121 | 2.194851239 | 0.343 | 0.018 | 5.18E-117 |
| **CTSA** | 4.47E-119 | 2.6297131 | 0.778 | 0.107 | 1.49E-114 |
| **AC127502.2** | 2.46E-112 | 2.295650198 | 0.404 | 0.027 | 8.23E-108 |
| **KIF2A** | 1.64E-109 | 1.946149459 | 0.374 | 0.023 | 5.48E-105 |
| **RGS18** | 5.01E-109 | 2.496340578 | 0.798 | 0.119 | 1.67E-104 |
| **ILK** | 1.40E-97 | 2.07872366 | 0.495 | 0.047 | 4.68E-93 |
| **MPP1** | 3.73E-95 | 2.389218665 | 0.657 | 0.088 | 1.25E-90 |
| **PLA2G12A** | 1.16E-94 | 1.864884362 | 0.414 | 0.034 | 3.87E-90 |
| **INAFM2** | 3.33E-94 | 1.783075682 | 0.323 | 0.02 | 1.11E-89 |
| **SMIM3** | 1.92E-91 | 1.823561482 | 0.343 | 0.024 | 6.43E-87 |
| **STON2** | 3.61E-90 | 1.79474786 | 0.263 | 0.014 | 1.21E-85 |
| **NCK2** | 5.71E-90 | 2.173935735 | 0.556 | 0.066 | 1.91E-85 |
| **STOM** | 2.22E-89 | 2.30691427 | 0.535 | 0.062 | 7.43E-85 |
| **MAX** | 5.29E-89 | 2.682610982 | 0.808 | 0.157 | 1.77E-84 |
| **NAP1L1** | 3.66E-84 | 2.42886834 | 0.859 | 0.203 | 1.22E-79 |
| **RNF11** | 7.29E-82 | 2.151391835 | 0.475 | 0.051 | 2.44E-77 |
| **SCN1B** | 9.68E-81 | 1.584954692 | 0.293 | 0.019 | 3.23E-76 |
| **PARVB** | 1.77E-80 | 1.746229301 | 0.313 | 0.022 | 5.92E-76 |
| **TUBA4A** | 3.02E-80 | 2.003782856 | 0.758 | 0.141 | 1.01E-75 |
| **CCL5** | 4.58E-80 | 2.175146298 | 0.808 | 0.173 | 1.53E-75 |
| **LIMS1** | 3.81E-75 | 2.552614085 | 0.778 | 0.178 | 1.27E-70 |
| **GSTO1** | 1.24E-74 | 2.356791492 | 0.505 | 0.065 | 4.16E-70 |
| **UBL4A** | 1.53E-74 | 1.901312041 | 0.313 | 0.024 | 5.12E-70 |
| **ANO6** | 2.40E-74 | 1.839927191 | 0.354 | 0.031 | 8.03E-70 |
| **PTPN18** | 1.54E-71 | 2.267354661 | 0.576 | 0.09 | 5.13E-67 |
| **TPM4** | 1.17E-69 | 2.497357368 | 0.727 | 0.161 | 3.91E-65 |
| **TUBA1C** | 2.97E-69 | 2.122263255 | 0.303 | 0.024 | 9.93E-65 |
| **TUBA4B** | 8.87E-69 | 1.808031994 | 0.475 | 0.059 | 2.96E-64 |

| | | | | | |
|---|---|---|---|---|---|
| TLN1 | 5.76E-68 | 2.485536285 | 0.909 | 0.315 | 1.93E-63 |
| MYLK | 1.29E-67 | 2.106136589 | 0.434 | 0.052 | 4.32E-63 |
| BCL2L1 | 6.64E-67 | 1.707269646 | 0.313 | 0.027 | 2.22E-62 |
| OST4 | 2.27E-66 | 2.329115029 | 0.737 | 0.173 | 7.57E-62 |
| FLNA | 9.37E-60 | 2.223430155 | 0.848 | 0.279 | 3.13E-55 |
| NORAD | 1.27E-59 | 2.406524211 | 0.636 | 0.135 | 4.24E-55 |
| FERMT3 | 4.20E-57 | 2.141343883 | 0.636 | 0.14 | 1.40E-52 |
| PRUNE1 | 6.57E-54 | 1.401653871 | 0.263 | 0.023 | 2.19E-49 |
| NT5C3A | 2.50E-51 | 1.929517335 | 0.444 | 0.069 | 8.36E-47 |
| RAB32 | 1.37E-50 | 1.814342831 | 0.404 | 0.058 | 4.57E-46 |
| EIF2AK1 | 1.23E-49 | 1.927886928 | 0.374 | 0.051 | 4.09E-45 |
| CCDC92 | 3.66E-49 | 1.553970178 | 0.273 | 0.027 | 1.22E-44 |
| HIST2H2BE | 3.38E-48 | 1.783999793 | 0.263 | 0.026 | 1.13E-43 |
| H2AFJ | 9.45E-45 | 1.687623659 | 0.273 | 0.03 | 3.16E-40 |
| GNAS | 2.08E-44 | 1.880992577 | 0.909 | 0.441 | 6.93E-40 |
| GFI1B | 4.93E-44 | 1.545800134 | 0.283 | 0.033 | 1.65E-39 |
| CD226 | 4.96E-44 | 1.965662526 | 0.545 | 0.12 | 1.66E-39 |
| CALM3 | 1.36E-43 | 1.934573923 | 0.687 | 0.208 | 4.53E-39 |
| NCOA4 | 1.11E-41 | 1.799658759 | 0.838 | 0.348 | 3.71E-37 |
| OAZ1 | 3.67E-41 | 1.469169593 | 0.949 | 0.692 | 1.23E-36 |
| PRDX6 | 3.33E-40 | 1.738345795 | 0.394 | 0.067 | 1.11E-35 |
| MGAT4B | 6.57E-40 | 1.547962648 | 0.313 | 0.044 | 2.20E-35 |
| PIP4K2A | 7.34E-40 | 1.898673661 | 0.495 | 0.107 | 2.45E-35 |
| MGLL | 7.72E-40 | 1.579418801 | 0.283 | 0.036 | 2.58E-35 |
| PKM | 2.16E-39 | 1.927279682 | 0.566 | 0.147 | 7.20E-35 |
| MAPRE2 | 6.06E-39 | 1.696075615 | 0.384 | 0.066 | 2.02E-34 |
| RSU1 | 1.84E-38 | 2.03949499 | 0.465 | 0.1 | 6.13E-34 |
| ARHGAP21 | 1.89E-38 | 1.550341799 | 0.263 | 0.032 | 6.32E-34 |
| SNN | 3.48E-38 | 1.605605807 | 0.434 | 0.085 | 1.16E-33 |
| RAP1B | 7.99E-37 | 2.045204307 | 0.737 | 0.307 | 2.67E-32 |
| TPST2 | 5.11E-36 | 1.869780812 | 0.465 | 0.105 | 1.71E-31 |
| PLEKHO1 | 2.84E-35 | 1.850992198 | 0.535 | 0.14 | 9.49E-31 |
| LDLRAP1 | 6.44E-35 | 1.588585286 | 0.323 | 0.053 | 2.15E-30 |
| CAPN1 | 7.44E-35 | 1.603183603 | 0.404 | 0.082 | 2.48E-30 |
| TAGLN2 | 1.13E-31 | 1.299090612 | 0.879 | 0.515 | 3.76E-27 |
| TGFB1 | 1.27E-31 | 1.353457352 | 0.263 | 0.038 | 4.26E-27 |
| SH3BGRL3 | 6.61E-31 | 1.421497502 | 0.879 | 0.615 | 2.21E-26 |
| ETFA | 1.04E-30 | 2.015780758 | 0.414 | 0.096 | 3.46E-26 |
| PYCR2 | 3.93E-30 | 2.046355733 | 0.323 | 0.06 | 1.31E-25 |
| MFSD1 | 1.18E-29 | 1.464409228 | 0.323 | 0.06 | 3.94E-25 |
| MTPN | 1.22E-29 | 1.566711481 | 0.636 | 0.237 | 4.08E-25 |
| DAPP1 | 4.29E-29 | 1.495306712 | 0.444 | 0.11 | 1.43E-24 |
| CDK2AP1 | 7.12E-29 | 1.839106607 | 0.273 | 0.045 | 2.38E-24 |
| TPM1 | 8.97E-29 | 1.554912474 | 0.364 | 0.077 | 3.00E-24 |
| TIMP1 | 1.17E-28 | 1.711305293 | 0.495 | 0.139 | 3.90E-24 |
| ADIPOR1 | 2.23E-27 | 1.504978717 | 0.444 | 0.113 | 7.46E-23 |
| GPX4 | 5.22E-27 | 1.675844208 | 0.404 | 0.1 | 1.75E-22 |
| CMIP | 1.85E-26 | 1.580517878 | 0.525 | 0.17 | 6.19E-22 |
| TMSB4X | 2.78E-26 | 0.978131639 | 0.99 | 0.956 | 9.29E-22 |
| MYL12A | 3.11E-26 | 1.402922331 | 0.758 | 0.424 | 1.04E-21 |
| RBBP6 | 1.28E-25 | 1.472108077 | 0.303 | 0.06 | 4.26E-21 |
| RAB11A | 2.41E-25 | 1.618144979 | 0.374 | 0.088 | 8.04E-21 |
| CYB5R3 | 3.84E-25 | 1.577452327 | 0.354 | 0.083 | 1.28E-20 |
| LAMTOR1 | 6.99E-25 | 1.569000755 | 0.485 | 0.151 | 2.34E-20 |
| ITGB1 | 9.59E-25 | 1.834311763 | 0.475 | 0.147 | 3.20E-20 |
| CARD19 | 5.32E-24 | 1.6004179 | 0.303 | 0.064 | 1.78E-19 |
| DYNLL1 | 1.62E-23 | 1.894571821 | 0.374 | 0.099 | 5.41E-19 |
| CCND3 | 2.10E-23 | 1.517518244 | 0.566 | 0.223 | 7.01E-19 |

| | | | | | |
|---|---|---|---|---|---|
| **CORO1C** | 6.43E-23 | 1.637224145 | 0.273 | 0.053 | 2.15E-18 |
| **ACTB** | 7.28E-23 | 0.873975388 | 0.949 | 0.948 | 2.43E-18 |
| **WBP2** | 1.56E-22 | 1.380916389 | 0.475 | 0.148 | 5.20E-18 |
| **TLK1** | 2.18E-22 | 1.429358263 | 0.263 | 0.051 | 7.29E-18 |
| **UQCRH** | 3.51E-22 | 1.591304703 | 0.364 | 0.096 | 1.17E-17 |
| **PTPN12** | 4.72E-22 | 1.538794093 | 0.444 | 0.136 | 1.58E-17 |
| **TMEM140** | 7.22E-22 | 1.194915239 | 0.333 | 0.077 | 2.41E-17 |
| **CD99** | 9.77E-22 | 1.722610003 | 0.475 | 0.168 | 3.26E-17 |
| **RAP2B** | 2.33E-21 | 1.960244081 | 0.343 | 0.09 | 7.77E-17 |
| **MYL6** | 2.32E-20 | 1.437114033 | 0.707 | 0.422 | 7.77E-16 |
| **SNRPN** | 1.52E-19 | 1.367042551 | 0.263 | 0.058 | 5.09E-15 |
| **MYH9** | 1.73E-19 | 1.063344301 | 0.788 | 0.476 | 5.78E-15 |
| **PTPRJ** | 2.99E-19 | 1.391329089 | 0.333 | 0.088 | 9.98E-15 |
| **SLC40A1** | 5.82E-19 | 1.452357187 | 0.313 | 0.08 | 1.95E-14 |
| **SPINT2** | 6.60E-19 | 1.39068667 | 0.303 | 0.076 | 2.20E-14 |
| **F11R** | 6.92E-19 | 1.282123261 | 0.283 | 0.065 | 2.31E-14 |
| **WDR1** | 2.49E-18 | 1.350658709 | 0.535 | 0.224 | 8.31E-14 |
| **PDCD10** | 5.29E-18 | 1.193164361 | 0.253 | 0.056 | 1.77E-13 |
| **APP** | 9.84E-17 | 1.500001931 | 0.253 | 0.061 | 3.29E-12 |
| **MT-RNR2** | 7.61E-16 | 0.548976703 | 1 | 0.991 | 2.54E-11 |
| **RAB37** | 9.31E-16 | 1.227461055 | 0.283 | 0.075 | 3.11E-11 |
| **SERF2** | 4.56E-15 | 1.205721369 | 0.667 | 0.398 | 1.52E-10 |
| **HACD4** | 5.56E-15 | 1.107391155 | 0.303 | 0.088 | 1.86E-10 |
| **STK24** | 1.08E-14 | 1.452417302 | 0.354 | 0.121 | 3.62E-10 |
| **KCTD20** | 1.64E-14 | 1.149672657 | 0.343 | 0.113 | 5.48E-10 |
| **GSN** | 4.35E-14 | 1.140510886 | 0.374 | 0.132 | 1.45E-09 |
| **ARPC1B** | 4.75E-14 | 1.151555859 | 0.727 | 0.491 | 1.59E-09 |
| **CDKN2D** | 6.06E-14 | 1.17265665 | 0.444 | 0.182 | 2.02E-09 |
| **CERS2** | 7.56E-14 | 1.086836162 | 0.273 | 0.079 | 2.53E-09 |
| **TMBIM1** | 9.08E-14 | 1.414207179 | 0.364 | 0.132 | 3.03E-09 |
| **AMD1** | 9.53E-14 | 0.93367395 | 0.253 | 0.067 | 3.18E-09 |
| **ABCC3** | 9.72E-14 | 1.136755346 | 0.283 | 0.083 | 3.25E-09 |
| **H3F3A** | 1.49E-13 | 0.480229171 | 1 | 0.91 | 4.98E-09 |
| **DIAPH1** | 2.19E-13 | 1.296370107 | 0.364 | 0.139 | 7.33E-09 |
| **HLA-E** | 4.45E-13 | 0.581564919 | 0.929 | 0.873 | 1.49E-08 |
| **ARF1** | 5.34E-13 | 1.059772198 | 0.556 | 0.288 | 1.78E-08 |
| **MT-ND2** | 8.32E-13 | 0.834603014 | 0.788 | 0.585 | 2.78E-08 |
| **RASA3** | 1.08E-12 | 1.090170185 | 0.253 | 0.074 | 3.62E-08 |
| **MT-ND1** | 1.51E-11 | 0.702157566 | 0.919 | 0.719 | 5.05E-07 |
| **TUBA1B** | 2.15E-11 | 1.083992742 | 0.354 | 0.14 | 7.20E-07 |
| **SH3BGRL** | 3.44E-11 | 1.260354367 | 0.343 | 0.137 | 1.15E-06 |
| **RAB1B** | 4.12E-11 | 1.363501213 | 0.273 | 0.093 | 1.38E-06 |
| **NFE2** | 4.49E-11 | 1.026775973 | 0.273 | 0.089 | 1.50E-06 |
| **MT-RNR1** | 1.03E-10 | 0.505575122 | 0.97 | 0.869 | 3.43E-06 |
| **ASAP1** | 2.12E-10 | 1.044977296 | 0.374 | 0.156 | 7.08E-06 |
| **PTP4A2** | 3.01E-10 | 1.138724012 | 0.556 | 0.321 | 1.01E-05 |
| **SNAP23** | 4.13E-10 | 1.269699284 | 0.323 | 0.129 | 1.38E-05 |
| **MT-ATP6** | 5.14E-10 | 0.4389576 | 0.949 | 0.666 | 1.72E-05 |
| **YWHAZ** | 7.24E-10 | 1.160847306 | 0.586 | 0.395 | 2.42E-05 |
| **GAPDH** | 7.84E-10 | 0.911871946 | 0.596 | 0.397 | 2.62E-05 |
| **MTRNR2L12** | 8.24E-10 | 0.524494868 | 0.929 | 0.918 | 2.75E-05 |
| **BIN2** | 9.50E-10 | 1.043763434 | 0.515 | 0.297 | 3.17E-05 |
| **AP2M1** | 1.09E-09 | 1.225693393 | 0.313 | 0.129 | 3.65E-05 |
| **SQSTM1** | 1.77E-09 | 1.100822749 | 0.364 | 0.165 | 5.91E-05 |
| **USF2** | 3.43E-09 | 1.230521904 | 0.273 | 0.106 | 0.00011466 |
| **EMC3** | 3.83E-09 | 1.155856229 | 0.293 | 0.12 | 0.000127829 |
| **PPP3R1** | 1.19E-08 | 1.05906159 | 0.263 | 0.098 | 0.000398224 |
| **MBNL1** | 2.81E-08 | 1.269094538 | 0.434 | 0.258 | 0.000938879 |

| | | | | | |
|---|---|---|---|---|---|
| **MT-ND4** | 3.71E-08 | 0.446146018 | 0.889 | 0.674 | 0.001240651 |
| **TRAPPC1** | 5.74E-08 | 0.914087669 | 0.313 | 0.144 | 0.00191848 |
| **HDGF** | 6.09E-08 | 1.282589003 | 0.253 | 0.102 | 0.002034889 |
| **YPEL5** | 1.61E-07 | 1.060325054 | 0.343 | 0.162 | 0.005378502 |
| **GNAQ** | 1.81E-07 | 1.047636853 | 0.333 | 0.163 | 0.006035512 |
| **ACTN1** | 2.84E-07 | 0.879268987 | 0.495 | 0.296 | 0.009499725 |
| **PPDPF** | 3.20E-07 | 1.24773103 | 0.283 | 0.135 | 0.010690265 |
| **CAPNS1** | 4.17E-07 | 1.176749742 | 0.343 | 0.182 | 0.013918725 |
| **TPM3** | 7.77E-07 | 0.804540135 | 0.515 | 0.341 | 0.025953645 |
| **HMGB1** | 8.44E-07 | 1.037822948 | 0.515 | 0.358 | 0.028194787 |
| **WIPF1** | 8.70E-07 | 0.673452468 | 0.576 | 0.42 | 0.029069594 |
| **RAB8A** | 9.14E-07 | 1.041794751 | 0.263 | 0.119 | 0.030542986 |
| **EIF4G2** | 9.26E-07 | 0.841629524 | 0.444 | 0.278 | 0.03093416 |
| **ZYX** | 9.62E-07 | 0.635998302 | 0.535 | 0.354 | 0.03214706 |
| **IQGAP2** | 1.12E-06 | 0.940625017 | 0.283 | 0.13 | 0.037277383 |
| **MT-CYB** | 1.25E-06 | 0.414391391 | 0.747 | 0.519 | 0.041668321 |
| **UCP2** | 1.35E-06 | 0.868621204 | 0.343 | 0.184 | 0.045211161 |
| **MT-CO1** | 2.15E-06 | 0.2932858 | 0.788 | 0.485 | 0.071716656 |
| **DNAJB6** | 2.52E-06 | 1.113417214 | 0.283 | 0.137 | 0.084036421 |
| **TPP1** | 2.72E-06 | 1.094925411 | 0.283 | 0.141 | 0.090790024 |
| **RASGRP2** | 4.98E-06 | 0.867556275 | 0.263 | 0.124 | 0.166369805 |
| **PTMA** | 5.97E-06 | 0.656295057 | 0.515 | 0.341 | 0.199379858 |
| **ACTG1** | 9.06E-06 | 0.752639879 | 0.636 | 0.536 | 0.302754324 |
| **R3HDM4** | 1.21E-05 | 0.884710774 | 0.293 | 0.146 | 0.403897135 |
| **PLEK** | 8.27E-05 | 0.982262127 | 0.404 | 0.275 | 1 |
| **PRKCB** | 0.000107441 | 0.734843552 | 0.485 | 0.343 | 1 |
| **PARK7** | 0.000123739 | 0.670701792 | 0.283 | 0.158 | 1 |
| **MOB1A** | 0.000129527 | 0.853245355 | 0.313 | 0.189 | 1 |
| **CAPZB** | 0.000150245 | 0.578603034 | 0.495 | 0.363 | 1 |
| **TMEM50A** | 0.00015275 | 0.692831921 | 0.283 | 0.157 | 1 |
| **SLC44A2** | 0.000178418 | 0.564757222 | 0.384 | 0.231 | 1 |
| **MT-ND5** | 0.000777686 | 0.534390941 | 0.556 | 0.415 | 1 |
| **MT-ND3** | 0.000935462 | 0.266242372 | 0.818 | 0.6 | 1 |
| **ARL6IP5** | 0.001071443 | 0.891812963 | 0.283 | 0.185 | 1 |
| **UBB** | 0.00327809 | 0.687974962 | 0.414 | 0.309 | 1 |
| **CAPZA2** | 0.004109203 | 0.611067512 | 0.364 | 0.257 | 1 |
| **FXYD5** | 0.005570768 | 1.098204351 | 0.273 | 0.19 | 1 |
| **SAT1** | 0.005575991 | 0.380539294 | 0.556 | 0.463 | 1 |
| **MTRNR2L8** | 0.008028275 | 0.413926193 | 0.283 | 0.187 | 1 |
| **FKBP1A** | 0.034585847 | 0.476781825 | 0.323 | 0.259 | 1 |
| **SEC14L1** | 0.057575064 | 0.535491374 | 0.303 | 0.235 | 1 |
| **SLC2A3** | 0.065240971 | 0.392133435 | 0.343 | 0.278 | 1 |
| **ARPC4** | 0.099997173 | 0.392592708 | 0.273 | 0.216 | 1 |
| **CD63** | 0.103594297 | 0.472802389 | 0.283 | 0.227 | 1 |
| **CALM1** | 0.164161309 | 0.444187353 | 0.354 | 0.327 | 1 |
| **RHOA** | 0.220113572 | 0.315134152 | 0.525 | 0.519 | 1 |
| **ATP6V0E1** | 0.260914847 | 0.276159359 | 0.253 | 0.217 | 1 |
| **HLA-A** | 0.261744897 | 0.294519538 | 0.747 | 0.765 | 1 |
| **MT-ND4L** | 0.321103868 | 0.633046995 | 0.283 | 0.262 | 1 |
| **FYB1** | 0.486413554 | 0.250851952 | 0.505 | 0.496 | 1 |
| **ASAH1** | 0.54267204 | 0.519426742 | 0.293 | 0.287 | 1 |
| **SRP14** | 0.82403964 | 0.46879532 | 0.222 | 0.26 | 1 |
| **PPIA** | 0.879535949 | 0.277686784 | 0.303 | 0.326 | 1 |

### Table S4. List of upregulated genes in Neu 5 as compared to all other leukocytes.

| Genes | p_val | avg_logFC | pct.1 | pct.2 | p_val_adj |
|---|---|---|---|---|---|
| PPBP | 0 | 1.456431005 | 0.32 | 0.015 | 0 |
| PF4 | 3.24E-294 | 1.136572498 | 0.281 | 0.014 | 1.08E-289 |
| NRGN | 6.33E-270 | 1.745662672 | 0.407 | 0.04 | 2.12E-265 |
| MT-RNR2 | 1.34E-48 | 0.392564749 | 0.995 | 0.991 | 4.48E-44 |
| FTH1 | 6.43E-37 | 0.413653788 | 0.986 | 0.899 | 2.15E-32 |
| NAMPT | 5.24E-30 | 0.420289641 | 0.883 | 0.61 | 1.75E-25 |
| FTL | 1.12E-28 | 0.391499551 | 0.913 | 0.863 | 3.74E-24 |
| NEAT1 | 5.09E-27 | 0.351545077 | 0.961 | 0.825 | 1.70E-22 |
| MTRNR2L12 | 2.51E-26 | 0.331944525 | 0.952 | 0.917 | 8.38E-22 |
| CXCR2 | 1.44E-25 | 0.411681156 | 0.84 | 0.614 | 4.81E-21 |
| MT-RNR1 | 5.12E-25 | 0.356856589 | 0.92 | 0.868 | 1.71E-20 |
| IFITM2 | 8.50E-25 | 0.337618002 | 0.979 | 0.874 | 2.84E-20 |
| FCGR3B | 8.14E-22 | 0.436339851 | 0.744 | 0.518 | 2.72E-17 |
| CSF3R | 3.71E-21 | 0.366931905 | 0.796 | 0.599 | 1.24E-16 |
| DUSP1 | 8.24E-20 | 0.437075974 | 0.696 | 0.516 | 2.75E-15 |
| RGS2 | 3.88E-18 | 0.35703297 | 0.789 | 0.603 | 1.30E-13 |
| OAZ1 | 7.73E-15 | 0.365824651 | 0.764 | 0.692 | 2.58E-10 |
| LITAF | 3.68E-13 | 0.282353979 | 0.815 | 0.736 | 1.23E-08 |
| SOD2 | 6.25E-13 | 0.269518966 | 0.741 | 0.593 | 2.09E-08 |
| MNDA | 1.16E-12 | 0.315805029 | 0.659 | 0.513 | 3.87E-08 |
| S100A11 | 1.98E-12 | 0.292649476 | 0.769 | 0.674 | 6.61E-08 |
| Metazoa-SRP | 8.66E-12 | 0.251960935 | 0.874 | 0.86 | 2.89E-07 |
| YPEL3 | 2.52E-11 | 0.472828242 | 0.508 | 0.396 | 8.41E-07 |
| FOS | 2.92E-11 | 0.459890387 | 0.412 | 0.279 | 9.76E-07 |
| TAGLN2 | 3.86E-11 | 0.37609061 | 0.622 | 0.514 | 1.29E-06 |
| NCF1 | 8.25E-11 | 0.443491485 | 0.513 | 0.382 | 2.76E-06 |
| HLA-E | 9.51E-11 | 0.271668192 | 0.867 | 0.874 | 3.18E-06 |
| FPR1 | 1.36E-10 | 0.370462284 | 0.602 | 0.483 | 4.54E-06 |
| RIPOR2 | 1.93E-10 | 0.301551534 | 0.703 | 0.652 | 6.46E-06 |
| RTN3 | 1.97E-10 | 0.50656653 | 0.366 | 0.253 | 6.57E-06 |
| SLC25A37 | 2.79E-10 | 0.302287954 | 0.622 | 0.477 | 9.33E-06 |
| IFITM3 | 7.12E-10 | 0.55991834 | 0.334 | 0.224 | 2.38E-05 |
| BASP1 | 2.33E-09 | 0.251407761 | 0.659 | 0.519 | 7.80E-05 |
| MMP25 | 7.05E-09 | 0.333072804 | 0.398 | 0.277 | 0.000235616 |
| GPSM3 | 1.74E-08 | 0.493546718 | 0.327 | 0.23 | 0.000582696 |
| H3F3B | 1.82E-08 | 0.254839625 | 0.748 | 0.713 | 0.000607439 |
| MBOAT7 | 2.01E-08 | 0.340922915 | 0.414 | 0.299 | 0.000672688 |
| RAB31 | 3.27E-08 | 0.488258882 | 0.407 | 0.309 | 0.00109393 |
| PHC2 | 3.29E-08 | 0.38745142 | 0.405 | 0.305 | 0.001099635 |
| LYN | 4.10E-08 | 0.32414819 | 0.593 | 0.5 | 0.001370946 |
| KCTD21-AS1 | 4.79E-08 | 0.383891537 | 0.54 | 0.466 | 0.001598785 |
| TSC22D3 | 5.32E-08 | 0.361937101 | 0.59 | 0.52 | 0.001777256 |
| EVI2B | 7.01E-08 | 0.309733194 | 0.595 | 0.511 | 0.002340802 |
| PTRH1 | 7.67E-08 | 0.432019408 | 0.336 | 0.247 | 0.002560986 |
| RNASET2 | 1.48E-07 | 0.43102506 | 0.503 | 0.434 | 0.004940566 |
| IDH2 | 1.83E-07 | 0.501701426 | 0.318 | 0.235 | 0.00610306 |
| S100P | 3.60E-07 | 0.336384717 | 0.295 | 0.197 | 0.012035432 |
| MXD1 | 3.85E-07 | 0.280991668 | 0.476 | 0.36 | 0.012878098 |
| USP10 | 4.71E-07 | 0.404599031 | 0.359 | 0.264 | 0.015727224 |
| AQP9 | 1.53E-06 | 0.307943545 | 0.352 | 0.253 | 0.051001099 |
| TALDO1 | 2.56E-06 | 0.299329787 | 0.535 | 0.449 | 0.08547837 |
| CXCR1 | 2.60E-06 | 0.387833027 | 0.405 | 0.313 | 0.086891918 |
| CEBPB | 4.40E-06 | 0.346692077 | 0.533 | 0.463 | 0.147128218 |
| LAPTM5 | 7.06E-06 | 0.255435434 | 0.696 | 0.67 | 0.235880168 |
| SAT1 | 8.37E-06 | 0.255972357 | 0.54 | 0.462 | 0.279758922 |
| MCL1 | 1.04E-05 | 0.29022391 | 0.622 | 0.59 | 0.347199685 |

| | | | | | |
|---|---|---|---|---|---|
| ABTB1 | 1.37E-05 | 0.444760042 | 0.295 | 0.219 | 0.457758744 |
| MSRB1 | 1.54E-05 | 0.334058437 | 0.444 | 0.358 | 0.51462849 |
| SDCBP | 1.75E-05 | 0.291851478 | 0.563 | 0.496 | 0.585653159 |
| ACTN1 | 1.79E-05 | 0.379369136 | 0.373 | 0.295 | 0.596477495 |
| IRS2 | 2.28E-05 | 0.438790305 | 0.325 | 0.26 | 0.762484506 |
| ICAM3 | 2.33E-05 | 0.313297135 | 0.378 | 0.301 | 0.780016589 |
| XPO6 | 2.62E-05 | 0.311209226 | 0.364 | 0.282 | 0.873950507 |
| CYP4F3 | 2.75E-05 | 0.382931855 | 0.284 | 0.206 | 0.919631535 |
| CDA | 2.97E-05 | 0.378405737 | 0.3 | 0.227 | 0.991555304 |
| SLC11A1 | 3.21E-05 | 0.369296246 | 0.327 | 0.257 | 1 |
| FMNL1 | 4.01E-05 | 0.329483792 | 0.426 | 0.375 | 1 |
| MYADM | 5.16E-05 | 0.343812674 | 0.357 | 0.291 | 1 |
| FFAR2 | 5.26E-05 | 0.374060496 | 0.304 | 0.232 | 1 |
| SEC14L1 | 6.75E-05 | 0.324746655 | 0.307 | 0.233 | 1 |
| UBALD2 | 0.000131278 | 0.295540155 | 0.38 | 0.32 | 1 |
| SMAP2 | 0.000134198 | 0.34440321 | 0.346 | 0.287 | 1 |
| ANP32A | 0.000196772 | 0.307211672 | 0.437 | 0.387 | 1 |
| NADK | 0.000320866 | 0.327674329 | 0.309 | 0.249 | 1 |
| STX3 | 0.000378616 | 0.327384512 | 0.263 | 0.202 | 1 |
| ACSL1 | 0.000404462 | 0.257544014 | 0.343 | 0.277 | 1 |
| CLEC4E | 0.000414663 | 0.324351111 | 0.261 | 0.201 | 1 |
| NINJ1 | 0.00043459 | 0.262532617 | 0.3 | 0.236 | 1 |
| SLAMF6 | 0.001063634 | 0.435250584 | 0.341 | 0.297 | 1 |
| LRRC25 | 0.001253033 | 0.524140052 | 0.263 | 0.217 | 1 |
| CPPED1 | 0.001286237 | 0.323802613 | 0.368 | 0.317 | 1 |
| OAZ2 | 0.001681322 | 0.261089733 | 0.339 | 0.285 | 1 |
| PTEN | 0.001789145 | 0.361467704 | 0.323 | 0.275 | 1 |
| UBE2B | 0.002838563 | 0.285562506 | 0.366 | 0.316 | 1 |
| SKAP2 | 0.002984842 | 0.286780314 | 0.309 | 0.262 | 1 |
| SERPINA1 | 0.003282041 | 0.292079465 | 0.348 | 0.306 | 1 |
| VASP | 0.003887014 | 0.267439537 | 0.343 | 0.299 | 1 |
| STAT3 | 0.004109277 | 0.331298767 | 0.295 | 0.259 | 1 |
| NCOA4 | 0.00476307 | 0.271021703 | 0.38 | 0.35 | 1 |
| LINC01001 | 0.006562419 | 0.27721254 | 0.302 | 0.258 | 1 |
| MSL1 | 0.00665961 | 0.271563409 | 0.307 | 0.265 | 1 |
| HELLPAR | 0.006931274 | 0.317172564 | 0.421 | 0.421 | 1 |
| ARHGAP9 | 0.011251766 | 0.276996123 | 0.311 | 0.275 | 1 |
| MARCKS | 0.021677688 | 0.340264894 | 0.261 | 0.225 | 1 |
| ADAM8 | 0.022625013 | 0.273663445 | 0.275 | 0.242 | 1 |
| VSIR | 0.034912501 | 0.27232263 | 0.382 | 0.367 | 1 |
| ZFP36 | 0.037945494 | 0.308765153 | 0.295 | 0.274 | 1 |
| LILRB3 | 0.043752911 | 0.270031295 | 0.314 | 0.29 | 1 |
| ANXA11 | 0.044492155 | 0.368121174 | 0.307 | 0.293 | 1 |
| ERGIC1 | 0.049119215 | 0.265982888 | 0.284 | 0.262 | 1 |
| RAP1A | 0.060279949 | 0.404246298 | 0.281 | 0.269 | 1 |
| MTPN | 0.111215763 | 0.406568245 | 0.252 | 0.24 | 1 |
| RNF130 | 0.111522752 | 0.251202623 | 0.256 | 0.237 | 1 |
| ZYX | 0.119821355 | 0.28764036 | 0.357 | 0.355 | 1 |
| TLN1 | 0.233302827 | 0.256271371 | 0.309 | 0.32 | 1 |
| PRKCB | 0.284784323 | 0.258349278 | 0.33 | 0.345 | 1 |
| GTF2I | 0.341855547 | 0.262483594 | 0.281 | 0.285 | 1 |
| IGF2R | 0.561626681 | 0.282583998 | 0.261 | 0.272 | 1 |
| CACNB4 | 0.64418298 | 0.260429032 | 0.261 | 0.283 | 1 |
| BAZ2B | 0.831413205 | 0.260794652 | 0.229 | 0.254 | 1 |

**Table S5. Platelet gene-free specific NPA signature as opposed to the remaining neutrophils clusters.**

| **120_gene_NPA_signature** |
|---|
| PADI4 |
| MMP9 |
| RFLNB |
| S100A12 |
| HNRNPA2B1 |
| RTN3 |
| PYGL |
| RPLP1 |
| ADD3 |
| MT-ATP8 |
| NXPE3 |
| MT-CO2 |
| CYLD |
| GRN |
| TSPO |
| GPR82 |
| CYSLTR1 |
| RAB3D |
| NFATC2IP |
| TXN |
| C2orf15 |
| ZNF106 |
| USP10 |
| IDH2 |
| CTSD |
| MDM4 |
| SUPT4H1 |
| MME |
| SPAG9 |
| FLOT2 |
| CRISPLD2 |
| IL10RB |
| YY1 |
| MIGA1 |
| CDA |
| GTF2I |
| ADAR |
| RPL27A |
| EIF1 |
| TRIM41 |
| TMUB2 |
| G6PD |
| CYP4F3 |
| B4GALT5 |
| RAB5B |
| VAPA |
| N4BP2L2 |
| CCNI |
| IFRD1 |
| AC135012.1 |
| PGD |
| HMGB2 |
| ZNF397 |
| PSMA3-AS1 |
| PTRH1 |
| GNAI2 |

| |
|---|
| HSPA1A |
| ALOX5AP |
| HELLPAR |
| MGAM |
| RPL34 |
| TPT1 |
| RBPJ |
| SPI1 |
| S100A9 |
| TSC22D4 |
| EEF1A1 |
| IRS2 |
| TSC22D3 |
| ADGRE2 |
| FTH1 |
| SLAMF6 |
| ATP6V1A |
| IFITM3 |
| MBOAT7 |
| ACAP2 |
| RAP1A |
| KIAA1328 |
| NEDD9 |
| CXCR2 |
| FFAR2 |
| MSN |
| TXNIP |
| PAK2 |
| RPL41 |
| SPATS2 |
| FAU |
| ANXA11 |
| C20orf24 |
| RAC2 |
| RGS19 |
| FOS |
| NMNAT1 |
| LILRA2 |
| MT-CO3 |
| VASP |
| ANP32A |
| S100P |
| C19orf47 |
| MAPK1 |
| CKLF |
| STK10 |
| TMBIM6 |
| LILRB3 |
| PCSK7 |
| SARAF |
| RBBP5 |
| TAOK3 |
| FKBP5 |
| ALOX5 |
| GPR155 |
| PLXNC1 |
| LRRC25 |
| STAG2 |

| |
|---|
| DDX17 |
| CFL1 |
| RAB27A |
| KLF2 |
| CSF2RB |
| SELPLG |

**Table S6. NPAs-associated neutrophil signature as opposed to the remaining neutrophils clusters and all the other leukocytes.**

| 16_gene_NPA_signature |
|---|
| RTN3 |
| RAB3D |
| IL10RB |
| GPR82 |
| SPAG9 |
| B4GALT5 |
| RBPJ |
| IDH2 |
| TMUB2 |
| ZNF397 |
| NXPE3 |
| TRIM41 |
| HELLPAR |
| CCNI |
| TSC22D4 |
| YY1 |
